# New sensitive tools to characterize meta-metabolome response to short- and long-term cobalt exposure in dynamic river biofilm communities

**DOI:** 10.1101/2023.11.16.567369

**Authors:** Simon Colas, Benjamin Marie, Soizic Morin, Mathieu Milhe-Poutingon, Pierre Foucault, Siann Chalvin, Clémentine Gelber, Patrick Baldoni-Andrey, Nicholas Gurieff, Claude Fortin, Séverine Le Faucheur

**Author notes:** Corresponding author at: Universite de Pau et des Pays de l’Adour, E2S-UPPA, CNRS, IPREM, Pau, France. *E-mail address* (S. Colas).

## Abstract

Untargeted metabolomics is a non-*a priori* analysis of biomolecules that characterizes the metabolome variations induced by short- and long-term exposures to stressors. Even if the metabolite annotation remains lacunar due to database gaps, the global metabolomic fingerprint allows for trend analyses of dose-response curves for hundreds of cellular metabolites. The combination of untargeted metabolomic features and benchmark-dose (BMD) calculations then makes it possible to determine concentration range inducing defense responses (CRIDeR) and concentration range inducing damage responses (CRIDaR). To develop this approach in a context of time-dependent microbial community changes, mature river biofilms were exposed for 1 month to four cobalt (Co) concentrations (background concentration, 1 x 10^-7^, 5 x 10^-7^ and 1 x 10^-6^ M) in an open system of artificial streams. The meta-metabolomic response of biofilms was compared against a multitude of biological parameters (including bioaccumulation, biomass, chlorophyll *a* content, composition and structure of prokaryotic and eukaryotic communities) monitored at set exposure times (from 1 hour to 28 days). Cobalt exposure induced extremely rapid responses of the meta-metabolome, with time range inducing defense responses (TRIDeR) of around ten seconds, and time range inducing damage responses (TRIDaR) of several hours. Even in biofilms whose structure had been altered by Co bioaccumulation (reduced biomass, chlorophyll *a* contents and changes in the composition and diversity of prokaryotic and eukaryotic communities), CRIDeRs with similar initiation thresholds (1.41 ± 0.77 x 10^-10^ M Co^2+^ added in the exposure medium) were set up at the meta-metabolome level at every time point. In contrast, the CRIDaR initiation thresholds increased by 10 times in long-term Co exposed biofilms. The present study demonstrates that defense and damage responses of biofilm meta-metabolome exposed to Co are rapidly and sustainably impacted, even within tolerant and resistant microbial communities.

**Graphical abstract:** 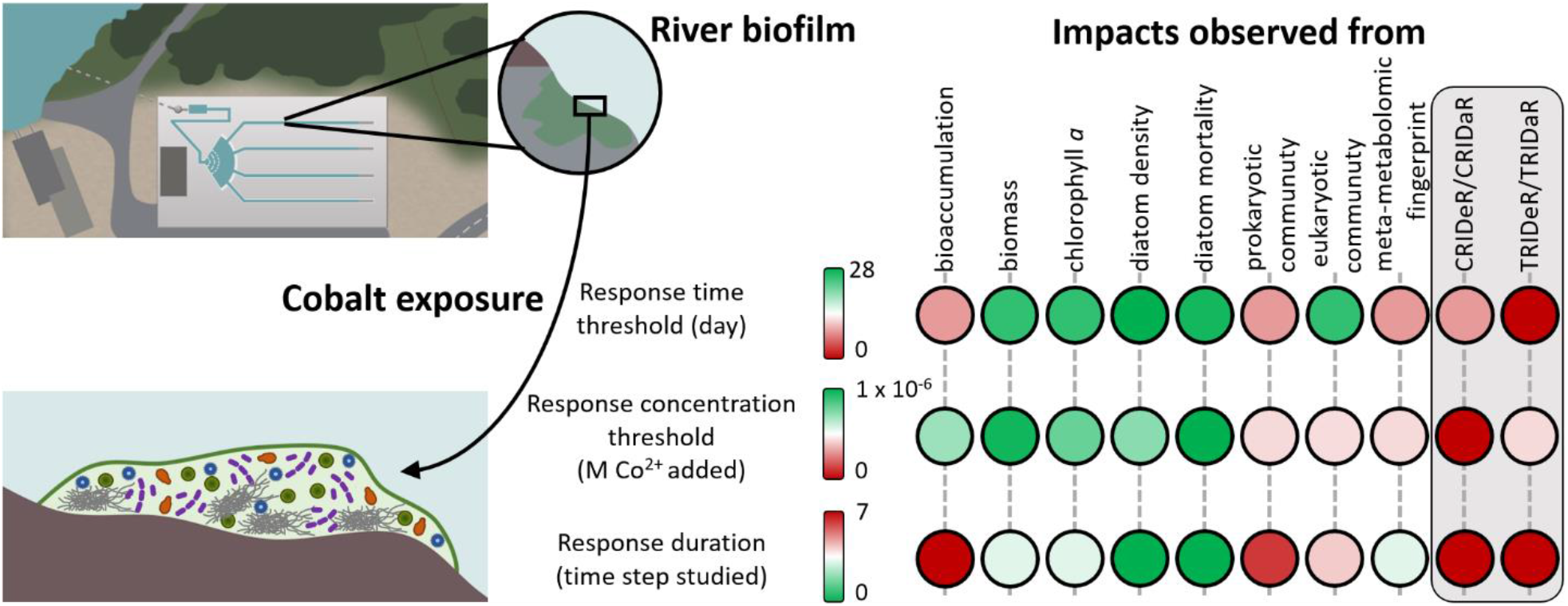

**Highlights:**

- Prokaryotic community structures were impacted after 1 h of exposure to Co
- Biofilm meta-metabolome was impacted after 36 s of exposure to Co
- Biofilm meta-metabolome response was faster than changes in biofilm communities
- Short- and long-term exposed biofilms have similar CRIDeR initiation thresholds
- Long-term exposed biofilms have higher CRIDaR initiation thresholds

## 1. Introduction

The current global transition towards decarbonized energy sources has led to the intensification of metal extraction. For example, cobalt (Co) mining has almost doubled since 2010, mostly due to its wide use in electric batteries (U.S. Geological Survey, 2010, 2023). As a consequence, increases in Co levels above background concentrations (< 10 nM) have been reported in aquatic ecosystems surrounding mining areas and industrial sites (Banza Lubaba Nkulu et al., 2018; Barrio-Parra et al., 2018). This metal is an essential trace element of biomolecules implicit in the metabolic pathways of various cellular processes (*e.g.*, vitamin B12; Martens et al., 2002). Nevertheless, experiments performed with model organisms, such as microalgae, demonstrated its toxic effects at µM concentrations on growth (Karthikeyan et al., 2019), lipid peroxidation (Li et al., 2007), pigment contents and photosynthetic activity (El-Sheekh et al., 2003; Plekhanov and Chemeris, 2003; Fawzy et al., 2020).

In rivers, benthic microalgae grow within biofilms, which are communities composed of additional types of organisms (*i.e.*, bacteria, fungi and meiofauna). They colonize various submerged substrata within aquatic systems and constitute the basis of the aquatic food web as primary producers. Biofilms are also well-known to accumulate metals as a function of their respective ambient concentration and speciation (Meylan et al., 2003; Lavoie et al., 2012a; Laderriere et al., 2020). Metals can then induce numerous effects at all levels of biological organization, from subcellular (Barranguet et al., 2003; Bonet et al., 2013), cellular (Morin et al., 2008b; Duong et al., 2010; Lavoie et al., 2012b; Morin et al., 2017) through to community (Barranguet et al., 2003; Bonet et al., 2013; Doose et al., 2021).

Among the various toxicological effects to biofilms that can be monitored, dysregulation of metabolite composition has been observed to be correlated with contaminant concentration and linked to stresses (Gauthier et al., 2020). Nevertheless, the study of such meta-metabolomic (metabolomics at the community level) responses from biofilms following a stress exposure (including anthropogenic pressures) remains a promising approach, despite its recent emergence (Booth et al., 2011; Gauthier et al., 2020). To this end, characterization of the whole meta-metabolome using untargeted Liquid Chromatography-High Resolution Mass Spectrometry (LC-HRMS) would allow for a robust characterization of the meta-metabolomic fingerprint without requiring an *a priori* selection on the form and function of molecules being analyzed. It may also support the generation of a new understanding on the function of such complex biological systems that microbial biofilms represent. However, due to gaps in current molecular databases, metabolite annotation levels remain still very limited (da Silva et al., 2015) and concern around 5% of the whole metabolomic fingerprint signal (Dias et al., 2016).

However, a recent study proposed to develop an innovative approach using the trends observed in dose-response curves of meta-metabolomic features and the determination of the benchmark dose-responses (BMDs) to overcome this pitfall (Colas et al., 2023). As such, that approach takes the full advantage of meta-metabolomic investigations for ecotoxicology purposes. In this work, river biofilms were exposed to Co for 7 days and untargeted meta-metabolomic features were analyzed at each tested concentration. The construction of dose-response curves allowed the calculation of a BMD for each untargeted metabolite feature for a total of 481 BMDs. A BMD corresponds to the concentration at which the response, *i.e.*, a benchmark-response (BMR) of an untargeted meta-metabolomic feature differs significantly from a control. These BMDs were further aggregated into an empirical cumulative density function (ECDF) according to their trends, *i.e.*, biphasic (bell- or U-shaped) and monotonic (decreasing or increasing). Indeed, a recent meta-analysis demonstrated that the trends of dose-response model curves for biomarkers depend on their biological functions. As such, based on 2,595 observations made on 18 phyla exposed to inorganic or organic contaminants, cellular defense responses have been shown to have predominantly biphasic trends, while damage responses have been found to have mostly monotonic trends (Colas and Le Faucheur, 2023). Specific response concentration ranges could be determined using trend-specific ECDFs and then provide the concentrations at which defense or damage responses were predominant, according to concentration ranges inducing defense responses (CRIDeR) and concentration ranges inducing damage responses (CRIDaR), respectively. In biofilms exposed to Co, the CRIDeR initiation threshold was 1.7 x 10^-9^ M of Co^2+^ added in the exposure medium and the CRIDaR initiation threshold was 2.7 x 10^-6^ M, evident from decreases in biomass and chlorophyll *a* content measured. Pursuing this integrative approach, it may now be possible to also characterize a time range inducing defense responses (TRIDeR) and a time range inducing damage responses (TRIDaR) by interpretation of dose-response curve trends and benchmark-time (BMT) calculation to provide a time dimension to the response of the meta-metabolome to acute or chronic stress.

In the present study, we aimed to evaluate the extent that CRIDeR and CRIDaR would be pertinent in the case of a chronic metal contamination. Indeed, one could expect that biofilms may develop meta-metabolomic responses to cope with chronic metal stress, and that different exposure conditions will favor the development of tolerant species while inhibiting the development of metal-sensitive species, leading to a progressive change in microbial community structure. We therefore expected CRIDeR and CRIDaR to vary according to the exposure time. To verify that hypothesis, mature river biofilms were exposed to four Co concentrations (background concentration, 1 x 10^-7^, 5 x 10^-7^ and 1 x 10^-6^ M) in artificial rivers for 1 month. The relevance of CRIDeR and CRIDaR characterization for a microbial community during chronic exposure was assessed in comparison with more traditional biological parameters (biomass, chlorophyll *a* contents and diatom density, mortality and characterization of the prokaryotic and eukaryotic communities) at set sampling time points (hour 1, days 1, 3, 7, 14, 21 and 28). In addition, we monitored changes in the kinetics of biofilm microbial community composition (prokaryotes and eukaryotes) to understand their influence and potential relation with the meta-metabolomic response over time, and to gain some insight into the relevance of subsequent TRIDeR and TRIDaR. Thus, analyzing the meta-metabolome of biofilms following chronic contaminant exposure could be an innovative and holistic approach to characterize the risk assessment of contaminants in ecosystems.

## 2. Material and methods

### 2.1. Biofilm colonization and exposure to Co in artificial streams

The experiment was carried out from September 15 to November 09, 2021, at the outdoor TotalEnergies facility (pilot rivers) in Lacq (PERL, France) (Cailleaud et al., 2019). Mature biofilms were naturally colonized on glass slides (10 x 20 cm) in plastic boxes (eight slides per box) for a period of 4-weeks after being placed in the nursery of an open circuit supplied by the *Gave de Pau* river (Tien et al., 2009). Twelve streams (length, 40 m; width, 0.5 m; depth, 0.5 m; flow rate, 7.5 m^3^·h^-1^) were randomly assigned a Co exposure concentration, with a total of three replicate streams for each of the four exposure conditions (background concentration in the *Gave de Pau* used as the control, 1 x 10^-7^, 5 x 10^-7^ and 1 x 10^-6^ M).

Co (Cobalt(II) chloride hexahydrate, 98%, Thermo Scientific Chemicals) solutions of 1.09 x 10^-2^, 5.43 x 10^-2^ and 1.09 x 10^-1^ M were added to streams one day prior to beginning exposures (day 0) to obtain stabilized nominal Co concentrations of 1 x 10^-7^, 5 x 10^-7^ and 1 x 10^-6^ M. Cobalt solutions were injected continuously (71 mL·h^-1^) by a piston pump and mechanically dispersed in the water by shearing valves (Netzsch, Nemo, Germany) throughout the experiment. At the commencement of exposures, an open box containing mature biofilm slides was placed in each stream, 15 m downstream of the injection pump. The upstream nursery constituted of a reservoir containing living organisms, allowing for the promotion of natural and continuous colonization within the streams during the 28 days of experimentation. One biofilm slide from each stream was collected at set time points (1 h; 1, 3, 7, 14, 21 and 28 d). Subsamples from every slide were taken to determine total biofilm Co content, intracellular biofilm Co content, chlorophyll *a* content, meta-metabolome, prokaryotic and eukaryotic community characterization, and diatom populations. Lugol solution (Sigma-Aldrich, Germany) was added to each sample for diatom population monitoring (final concentration of 0.01% Lugol), which were then stored in the dark at 4°C. Collected biofilms to be analyzed for Co intracellular accumulation were placed in the dark at 4°C until treatment up to 3 hours (see section 2.3). Other samples were stored in the dark at - 20°C before lyophilization and further analyses.

### 2.2. Water analysis and Co speciation

Temperature, pH, conductivity, saturation and dissolved oxygen were measured using a multi-parameter field probe (Hach, IA, USA) just before biofilm sampling. Three samples (10 mL) of water were filtered through 0.45 μm cellulose acetate (CA) syringe filters (Ministart® NML, Startorius, Germany) and collected in polypropylene (PP) tubes (MetalFree®, Labcon, CA, USA) in each stream just before Co injection at each time point to determine dissolved metal concentrations. Three samples (10 mL) were additionally filtered on 0.45 µm CA filters and collected in PP tubes for cation and anion analyses with an additional two 50 mL samples filtered on 0.7 μm GF/F filters to be collected in burnt-amber glass flasks for dissolved organic carbon (DOC) concentrations at day 1, 7, 14, 21 and 28 in each stream. Cobalt concentrations in water were analyzed by Inductively Coupled Plasma Mass Spectrometry (ICP-MS model 7500, Agilent, CA, USA). The cation concentrations were measured by ICP-OES (Thermo iCAP 6000 Series, Thermo-Fisher, MA, USA) and the anion concentrations were measured by ion chromatography (Dionex Aquion, Thermo-Fisher, MA, USA). SRLS-6 certified material (River water, CNRC) was used to calibrate the accuracy of the ICP-MS analyses (recovery of 103 ± 5% for Co; n = 5) and the ICP-OES analyses (recovery of 98 ± 1% for Ca^2+^; n = 3). The DOC concentrations were measured by a ShimatzuTOC-L analyzer (Japan). For modeling Co speciation, the Windermere Humic Aqueous Model (WHAM) model VII with its default data base was used (Bryan et al., 2002; Tipping, 2007). We assumed that 50% of the dissolved organic matter (DOM) is made of carbon and that 65% of the DOM is fulvic acid, the remainder was considered as inert.

### 2.3. Determination of Co bioaccumulation

The same protocol was followed to determine total and intracellular Co concentrations in biofilm samples as in Colas et al. (2023). To the analysis of intracellular Co concentrations, fresh biofilm samples were rinsed with 10 mM of ethylenediaminetetraacetic acid (EDTA, in disodium salt dihydrate form, Sigma-Aldrich, Germany) for 10 min to remove adsorbed metals on the surface of the biofilm whereas no rinsing step was performed for analyzing total Co concentrations (Meylan et al., 2003; Olguín and Sánchez-Galván, 2012). After that, samples were lyophilized and digested with 70% HNO_3_ and 30% H_2_O_2_ prior to mineralization (UltraWAVE™ oven, Milestone, Italy). Total and intracellular metal concentrations were analyzed by ICP-MS (ICP-MS model 7500, Agilent, CA, USA). BCR-414 certified reference material (plankton, JRC, Brussels) was used to confirm the sample digestion and the analytical accuracy (recovery of 83 ± 5% for Co; n = 9). SLRS-6 was also used for this method to assess its accuracy (recovery of 108 ± 4% for Co, n = 5).

### 2.4. Biofilm structure characterization

The quantity of dry biomass (in g_DW_) per glass slide area (cm^2^) was obtained from biofilm samples collected for Co bioaccumulation, chorophyll *a* content, community structure and meta-metabolome analyses. Chlorophyll *a* contents were extracted with 10 mL of 90% acetone (Uvasol®, Sigma-Aldrich, Germany), analyzed by spectrophotometry (Lambda 750, PerkinElmer, MA, USA) and quantified following the Jeffrey and Humphrey protocol (Jeffrey and Humphrey, 1975). Diatom analyses were carried out at the XPO platform (INRAe, Cestas, France; doi: 10.17180/brey-mr38) (INRAe institut, Cestas, France). Diatom cell density and mortality were determined from Lugol-preserved material for all replicate samples using a Nageotte counting chamber (Morin et al., 2010), under light microscopy (*Olympus BX51* upright microscope, 400x magnification). Diatom counts were reported as cells·cm^-2^ of scraped substrate. The diatom replicate samples collected on day 28 were pooled to form one composite sample representative of the assemblage present for each Co condition. They were prepared according to the European standard EN ISO 13946 and counted using a Leica photomicroscope (oil immersion, 1000x magnification). A minimum of 400 individuals per sample (EN ISO 14407) were identified (Coste and Rosebery, 2011), with relative species abundance expressed as a %.

Subsequently to methanol extraction of metabolites, (described below in section 2.5), DNA was extracted from the pellets of the same samples (Duperron et al., 2023) using the QIAGEN DNeasy® PowerSoil® Pro Kit DNA extraction kit following the manufacturer’s instruction. The V4-V5 region of the 16S rRNA encoding gene was amplified using primers 515F (5’-GTGYCAGCMGCCGCGGTA-3’) and 928R (5’-ACTYAAAKGAATTGRCGGGG-3’)(Wang and Qian, 2009). The V8-V9 region of the 18S rRNA encoding gene was amplified using primers V8F (5’-ATAACAGGTCTGTGATGCCCT-3’) and 1510R (5’-CCTTCYGCAGGTTCACCTAC-3’)(Bradley et al., 2016). The amplicons were sequenced on an Illumina NextSeq 2000 300×2 bp platform (PGTB platform, Cestas, France; doi: 10.15454/1.5572396583599417E12). Sequence analyses were performed on the Galaxy FROGS pipeline (Afgan et al., 2018). Forward and reverse reads were timed at 280 and 275 bp, respectively. Amplicon Sequence Variants (ASVs) were obtained with the default parameters and affiliated with the 16S SILVA 138.1 AND 18S SILVA 138.1 databases for prokaryote and eukaryote community characterization, respectively. Alpha- and beta-diversity metrics were computed in RStudio (v4.1.2)(R Core Team, 2021) with the R package phyloseq (v1.38.0) (McMurdie and Holmes, 2013) and pairwiseAdonis (v0.4) (Arbizu, 2022). Principal coordinates analyses (PCoA) based on weighted and unweighted UniFrac distances were performed to explore the prokaryotic and eukaryotic phylogenetic dissimilarity within groups (combination of exposure time and Co concentration). Inter-group and intra-group variance were computed with PERMANOVA (999 permutations) and PERMDISP (999 permutations), respectively.

### 2.5. Meta-metabolomic analyses

As detailed previously in Colas et al. (2023), untargeted meta-metabolomic analyses were performed at the PtSMB plateform (Muséum National d’Histoire Naturelle, Paris, France) by LC-HRMS. From lyophilized biofilm samples (1 mg), metabolites were extracted with a solvent composed of 10 µL cold 75% methanol that was acidified with 0.1% formic acid. Biofilm samples were then sonicated on ice (SONICS Vibra Cell, Newton, CT, USA; 130 Watts, 20 kHz; 80% amplitude; 30 s) and centrifuged at 4°C (12,000 g; 10 min) (Colas et al., 2020). Supernatants were harvested and stored in the dark at −20°C.

For the mass spectrometry analysis, 2 µL of the supernatants containing the metabolite extracts were analyzed by Ultra-high-performance liquid chromatography (UH-PLC) (ELUTE, Bruker, Bremen, Germany) using a Polar Advance II 2.5 pore C18 (Thermo Fisher Scientific, Waltham, MA, USA) chromatographic column (300 µL·min^-1^ over a linear gradient of acidified (0.1% formic acid) acetonitrile from 5 to 90% in 15 min) coupled with a high-resolution mass spectrometer (ESI-QqTOF, Compact, Brucker, Bremen, Germany) at 2 Hz frequency in positive MS mode in the range of 50-1500 m/z. The software MetaboScape 4.0 (Bruker, Bremen, Germany) was used to recalibrate the MS data. Ions with intensities above 5,000 points in at least 10% of all samples were selected. AutoMS/MS analysis was then conducted in positive mode for the qualitative study of metabolites. A molecular network was constructed with MetGem 1.3.6. and the comparison of fragmentation profiles and metabolite annotation was conducted.

#### 2.5.1. Concentration range inducing defense and damage responses (CRIDeR/CRIDaR)

Untargeted meta-metabolomic features dose-response models were constructed using RStudio (v4.1.2) with the DRomics package (v2.5-0) (Larras et al., 2018). The biofilm meta-metabolomic response to Co^2+^ concentrations added in the exposure medium was determined at each time point. Half-minimum method for missing values was applied on the original data (peak intensity MS analysis) before log_2_ transformation. Significantly responding untargeted meta-metabolomic features to Co concentrations were then selected using an ANOVA with a false discovery rate (FDR) of 0.05. Dose-response models with the best AICc (second-order Akaike’s information criterion) were constructed. BMD_-1SD_ were calculated from a BMR_-1SD_ (SD is the residual standard deviation of the considered model). The BMD_-1SD_ represents the Co^2+^ concentration added in the exposure medium at which the response of each untargeted meta-metabolomic feature of exposed biofilms previously selected, differs from the response observed in control biofilms. Empirical cumulative density functions based on dose-response model trends (biphasic: bell- or U-shaped and monotonic: decreasing or increasing) were built in order to obtain an integrative response of biofilm untargeted meta-metabolomic features following Co exposure (Colas et al., 2023). On this basis, CRIDeR and CRIDaR initiation thresholds were determined for each time point from the best (with lowest AICc) fit model (*e.g.* Burr III, Gamma, Gompertz, Inverse-Pareto, Gumbel, log-logistic, log-logistic/log-logistic mix, log-normal, log-normal/log-normal mix and Weibull distributions) for ECDFs of untargeted meta-metabolomic features with biphasic (bell- or U-shaped) and monotonic (decreasing or increasing) trends, respectively, with the R package ssdtools (v1.0.6) (Thorley and Schwarz, 2018). The CRIDeR and CRIDaR initiation thresholds are the hazardous concentrations at which 5% of untargeted meta-metabolomic features with a biphasic or a monotonic trend, respectively, will potentially be affected on the basis of their BMDs. If all distributions failed to fit, the CRIDeR and CRIDaR initiation thresholds were determined directly with the BMD of the first untargeted meta-metabolomic feature impacted. The CRIDeRs end when the CRIDaRs begin.

#### 2.5.2. Time range inducing defense and damage responses (TRIDeR/TRIDaR)

Applying the same principle as CRIDeR/CRIDaR, untargeted meta-metabolomic features time-response models were built based on the biofilm meta-metabolomic response to Co exposure time for each exposure condition. After the log_2_ transformation step, significantly responding untargeted meta-metabolomic features to Co exposure time were selected using an ANOVA with an FDR of 0.05. Dose-response models with the best AICc were also constructed. BMT_-1SD_ were calculated from a BMR_-1SD_. The BMT_-1SD_ represents the exposure time (in days; log_10_ transformed) at which the response of each untargeted meta-metabolomic feature of exposed biofilms previously selected differs from the response observed in control biofilms. Empirical cumulative density function construction and TRIDeR and TRIDaR characterization were performed in the same way as for CRIDeR and CRIDaR (see section 2.5.1.). The TRIDeR and TRIDaR initiation thresholds are the hazardous exposure time at which 5% of untargeted meta-metabolomic features with a biphasic or a monotonic trend, respectively, will potentially be affected on the basis of their BMTs. If all distributions failed to fit, TRIDeR and TRIDaR initiation thresholds were determined directly with the BMT of the first untargeted meta-metabolomic feature impacted. The TRIDeRs end when the TRIDaRs begin.

### 2.6. Multivariate Omics Trajectory Analysis (MOTA)

The dynamics of biofilm meta-metabolome changes versus compositions of prokaryotic and eukaryotic biofilm communities were investigated by comparing trajectories of centroids using the Multivariate Omics Trajectory Analysis method (MOTA) (Foucault et al., 2022). Datasets of meta-metabolome, prokaryotic and eukaryotic biofilm communities were log_2_ transformed and analyzed identically. Median distances between Principal Component Analyses’ (PCA) centroid coordinates were computed by median to create a trajectory between each sampling time point for the four exposure conditions and displayed as the fraction of the total length achieved at each time point (from 0 to 100% between hour 1 to day 28). Trajectories were plotted for the exposure time (log_10_ transformed) versus meta-metabolomic data, 16S and 18S rRNA data for each exposure concentration, allowing comparison of their dynamics over time.

## 3. Results

### 3.1. Exposure conditions and media

The *Gave de Pau* water that flowed into the streams throughout the experiment had a pH of 8.29 ± 0.02 (Table A.1). Water temperature decreased during the exposure experiments, from 13.67 ± 0.31 °C at the start of exposure (day 0) to 10.28 ± 0.04 °C after 28 days (Table A.1). The average background total dissolved Co concentration for 28 days was 4.8 ± 3.5 x 10^-9^ M Co (1.36 ± 0.85 x 10^-9^ M Co^2+^), 8.32 ± 0.06 x 10^-4^ M Ca, 1.54 ± 0.02 x 10^-4^ M Mg, 0.40 ± 0.13 mg DOC·L^-1^ (Table A.1). The averages of total dissolved Co concentrations for 28 days for each nominal condition (1 x 10^-7^, 5 x 10^-7^ and 1 x 10^-6^ M) were 0.92 ± 0.07 x 10^-7^, 5.26 ± 0.35 x 10^-7^ and 1.09 ± 0.03 x 10^-6^ M, respectively (Table A.1). The corresponding average Co^2+^ concentrations were 2.68 ± 0.19 x 10^-8^, 1.68 ± 0.16 x 10^-7^ and 3.06 ± 0.34 x 10^-7^ M, respectively.

### 3.2. Metal bioaccumulation

Mature biofilm naturally contained similar levels of total accumulated Co and intracellular Co, indicating that Co contents were almost entirely intracellular with an average intracellular Co of 2.19 ± 0.20 x 10^-7^ mol·g_DW_^-1^. This natural intracellular Co concentration remained constant throughout the experiment (Fig. 1 and Table A.2). After 1 hour, total and intracellular concentrations at all other time points significantly increased with exposure concentrations (*post-hoc* Dunn test, *p* < 0.05; Fig. 1, A.1 and Table A.2). A bioaccumulation plateau was reached at day 14 for exposed biofilms (Fig. 1 and Table A.2). At the end of exposure (day 28), the majority of accumulated Co by biofilms was intracellular (71 ± 14%; n = 12). Cobalt exposure concentrations had no effect on the biofilm bioaccumulation of other metals (Li, Al, Mn, Fe, Cu, Zn, As, Pb, Cd and Pb) present in the *Gave de Pau* (Table A.3).

**Fig. 1:**
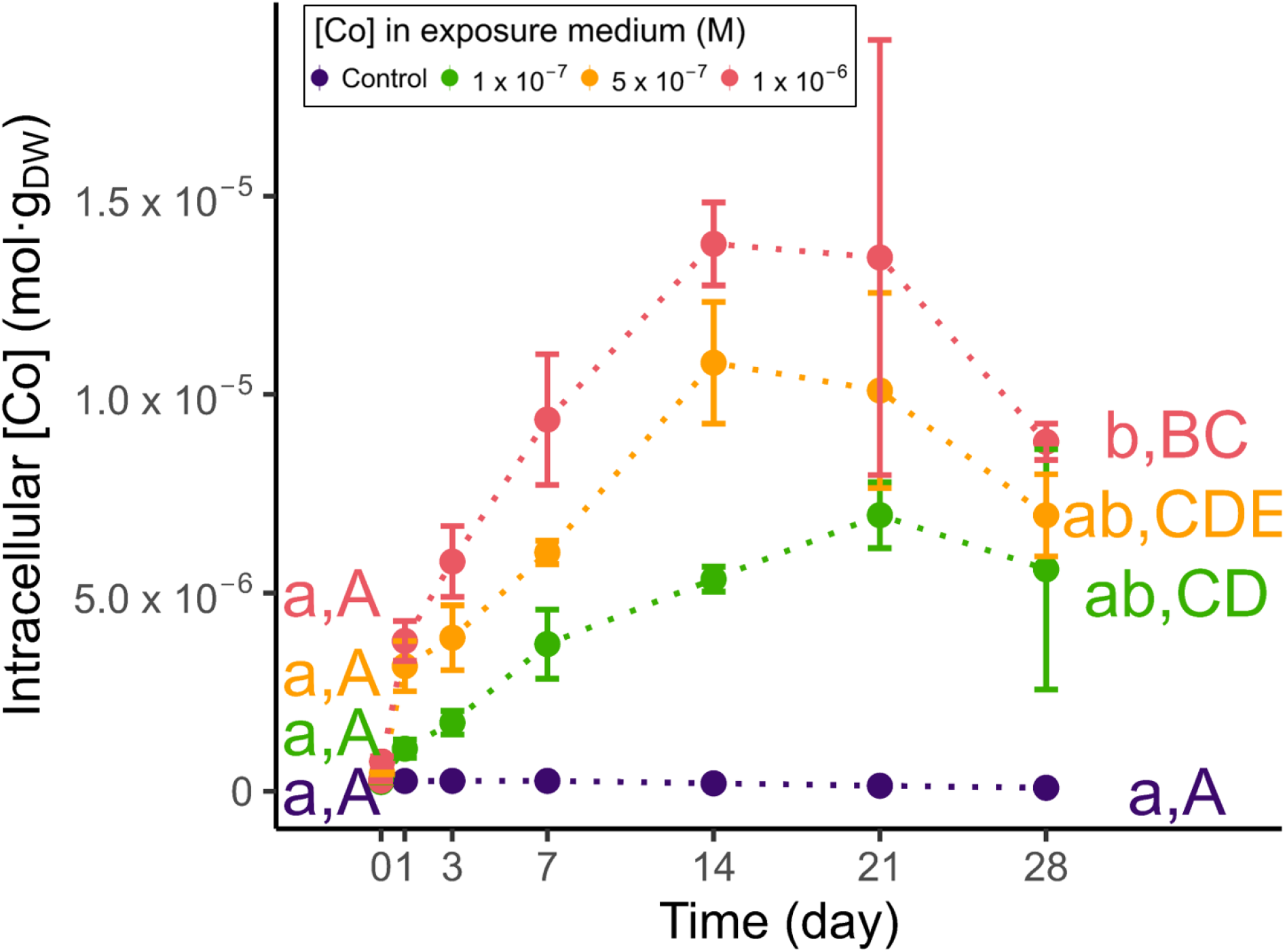
Intracellular Co content (mol·g^-1^_DW_) over time (day) according to Co concentration exposure groups (Control in blue, 1 x 10^-7^ M in green, 5 x 10^-7^ M in orange and 1 x 10^-6^ M in red). The lower-case letters correspond to the significative groups defined by a Dunn’s *post-hoc* test (*p* < 0.05) performed after Kruskal-Wallis non-parametric tests among Co exposure concentrations at each exposure time. Capital letters correspond to the significative groups defined by a Dunn’s *post-hoc* test (*p* < 0.05) performed after Kruskal-Wallis non-parametric tests among the exposure times for each exposure concentration. To improve make the graph easier to readability, only test results for days 0 and day 28 are displayed. Significance of tests at intermediate time points are displayed in Table A.2. (color should be used)

### 3.3. Biofilm structure

#### 3.3.1 Biomass and chlorophyll content

Biomass of control biofilms remained relatively stable over the 28-day experiment, with a slight but significant increase at days 3 and 7 (*post-hoc* Dunn test, *p* < 0.05, Table A.3). In contrast, biofilm biomass was negatively impacted by Co exposure from day 14 (Fig. 2A, 2B and Table A.4). On day 28, biofilm biomass decreased from 142 ± 12 mg_DW_·cm^-2^ in control biofilms to 17.8 ± 3.8 mg_DW_·cm^-2^ (−57%) and 17.8 ± 8.1 (−58%) mg_DW_·cm^-2^ at the two highest exposure concentrations (5 x 10^-7^ and 1 x 10^-6^ M), respectively (*post-hoc* Dunn test, *p* < 0.05). No effect was observed at 1 x 10^-7^ M Co at any exposure time.

**Fig. 2:**
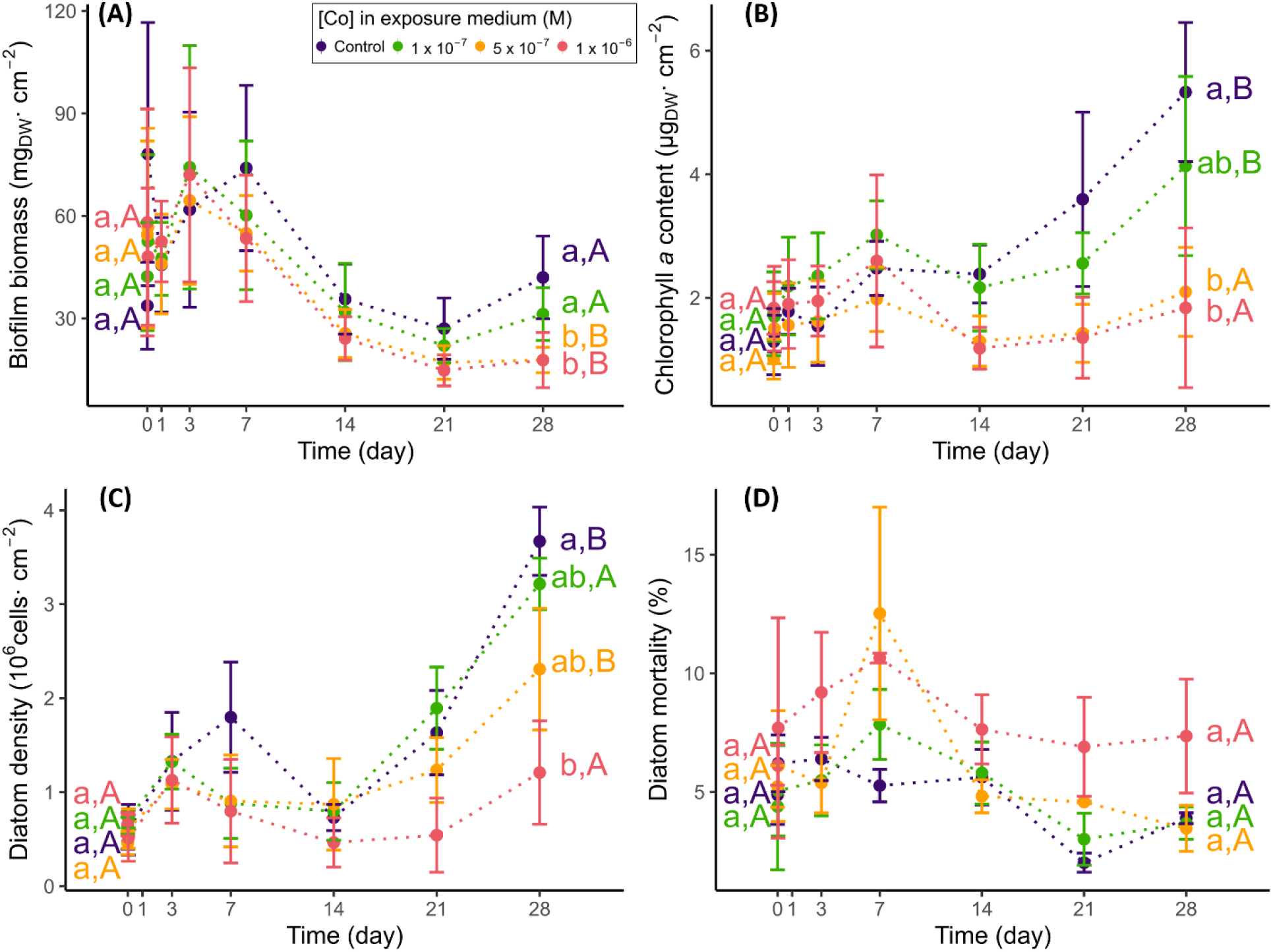
Effects of Co on traditional biological parameters of river biofilms (Control in blue, 1 x 10^-7^ M in green, 5 x 10^-7^ M in orange and 1 x 10^-6^ M in red). (A) Biofilm biomass (mg_DW_·cm^-2^) as a function of time (day) according to Co exposure concentrations. (B) Biofilm chlorophyll *a* content (μg_DW_·cm^-2^) as a function of time (day) according to Co exposure concentrations. (C) Diatom density (10^6^ cells·cm^-2^) as a function of time (day) according to Co exposure concentrations. (D) Diatom mortality (% of death cells) as a function of time (day) according to Co exposure concentrations. The lower-case letters correspond to the significative groups defined by a Dunn’s *post-hoc* test (*p* < 0.05) performed after Kruskal-Wallis non-parametric tests among Co exposure concentrations at each exposure time. Capital letters correspond to the significative groups defined by a Dunn’s *post-hoc* test (*p* < 0.05) performed after Kruskal-Wallis non-parametric tests among the exposure times for each exposure concentration. Only test results for days 0 and 28 are displayed, to make the graph easier to read. Significance of tests at intermediate time points are displayed in Table A.4. (color should be used)

Chlorophyll *a* content increased by a factor of 4- and 2.4-fold during the 28-day experiment in control biofilms and biofilms exposed to 1 x 10^-7^ M Co, respectively. At 5 x 10^-7^ and 1 x 10^-6^ M Co, chlorophyll *a* content decreased from 5.3 ± 1.1 in control biofilms to 2.10 ± 0.72 (− 61%) and 1.8 ± 1.3 (−65%) µg_DW_·cm^-2^, respectively (*post-hoc* Dunn test, *p* < 0.05; Fig. 2B and Table A.4).

#### 3.3.2. Prokaryotic biofilm community analysis

Analysis of prokaryotic community compositions based on the V4-V5 region of the 16S rRNA-encoding gene from the 192 biofilm metabolite-extract samples yielded 13,903,832 raw reads. Of which, 90.6% were retained after sequence assembly, denoising and chimera removal, and 74.1% were retained after a clustering filter (cluster with at least 0.005% of all sequences). Samples displayed from 41,886 to 121,693 reads (mean 69,951 ± 13,765 reads). Finally, 2,012 prokaryotic ASVs were taxonomically assigned.

At the first sampling time (1 h), the ASVs richness of control biofilms was 1866 ± 15 ASVs (Fig. A.2A). It remained constant up to 21 d of exposure, whereby it later significantly decreased by 13.3%; with comparable richness values found at days 21 and 28. Similar effects of exposure time were observed in Co-exposed biofilms. The ASVs richness decrease was exacerbated by Co at 28 d of exposure. It significantly decreased by 28.1%, 27.0% and 32.3% when compared to control biofilms at 1 x 10^-7^ M, 5 x 10^-7^ M and 1 x 10^-6^ M, respectively (*post-hoc* Dunn test, *p* < 0.05). The decreased ASVs richness of control biofilms at days 21 and 28 were accompanied by a significant 10% decrease in evenness (Fig. A.2B). In biofilms exposed to 1 x 10^-7^ M and 5 x 10^-7^ M Co, a significant decrease in evenness was observed at day 14, whereas it was only significant for biofilms exposed to the highest Co concentration at days 21 and 28. This decreased evenness was higher in biofilms exposed to 1 x 10^-6^ M Co for 28 d, with a value of 0.67 ± 0.02 as compared to the value of 0.75 ± 0.02 observed for control biofilms (*post-hoc* Dunn test, *p* < 0.05). The Shannon index also significantly decreased in control biofilms after 21 and 28 d of exposure (Fig. A.2C); a significant decrease from 6.34 to 5.66 and 5.56 were measured at days 21 and 28, respectively. With the Co exposures, a larger significant decrease was observed, with Shannon index decreases to 5.03, 4.93 and 4.76 in biofilms exposed to 1 x 10^-7^ M, 5 x 10^-7^ M and 1 x 10^-6^ M, respectively, on day 28 (*post-hoc* Dunn test, *p* < 0.05).

Control prokaryotic communities analyzed after 1 h of exposure consisted of over 16 identified classes, with a predominance of Cyanobacteria (27.9 ± 0.1%), Bacteroidia (14.8 ± 0.0%), Planctomycetes (13.3 ± 0.0%), Gammaproteobacteria (12.5 ± 0.0%) and Alphaproteobacteria (7.5± 0.0%) (Fig. A.3A). Visually, samples from different time points were well dispersed along the first axis of the PCoA, and exposure concentrations were rather separated on the second axis of the PCoA (Fig. 3A and 3B). At that early exposure time (1 h), the compositions of prokaryotic communities exposed to 5 x 10^-7^ and 1 x 10^-6^ M Co were already significantly different from those of the control biofilms (Unweighted UniFrac; PERMANOVA, *p* < 0.05; Fig. 3A; Table A.5). A significant decrease in the relative abundance of vadinHA49 (Planctomycetota) (*post-hoc* Dunn test, *p* < 0.05) and an increase in the relative abundance of Alphaproteobacteria *(post-hoc* Dunn test, *p* < 0.05) in biofilms exposed to 1 x 10^-6^ M were observed compared to those in control biofilms (Fig. A.3A). After 1 d of exposure, prokaryotic community compositions were significantly different only between those of control biofilms and those exposed to 5 x 10^-7^ M (Unweighted UniFrac; PERMANOVA, *p* < 0.05; Fig. 3A; Table A.5). On day 3, no significant differences were observed. On day 7, prokaryotic community compositions and diversities were significantly different only between control biofilms and biofilms exposed to 1 x 10^-6^ M (Unweighted UniFrac and Weighted UniFrac; PERMANOVA, *p* < 0.05; Fig. 3A and 3B; Table A.5). From day 14 to the end of the experiment (day 28), prokaryotic community compositions differed significantly between biofilms from the control group and biofilms exposed to 5 x 10^-7^ and 1 x 10^-6^ M (Unweighted UniFrac; PERMANOVA, *p* < 0.05; Fig. 3A; Table A.5). At days 21 and 28, prokaryotic community diversities differed significantly between biofilms from the control group and biofilms exposed to 5 x 10^-7^ and 1 x 10^-6^ M (Weighted UniFrac; PERMANOVA, *p* < 0.05; Fig. 3B; Table A.5). Prokaryotic community diversities were also significantly different in biofilms exposed to 1 x 10^-7^ and 1 x 10^-6^ M, and in biofilms exposed to 5 x 10^-7^ and 1 x 10^-6^ M (Weighted UniFrac; PERMANOVA, *p* < 0.05; Fig. 3B; Table A.5). At day 28, an increase in the relative abundance of the dominant Cyanobacteria class as a function of Co concentrations was observed, as well as a decrease of Polyangia, vadinHA49, KD4-96 (Chloroflexi) and OM190 (Planctomycetota) (*post-hoc* Dunn test, *p* < 0.05; Fig. A.3A). The diversities of prokaryotic communities of biofilms exposed to 1 x 10^-7^ M were never different from those of biofilms exposed to 5 x 10^-7^ M throughout the experiment whereas they were significantly different from those of biofilms exposed to 1 x 10^-6^ M on days 14 and 28 (Unweighted UniFrac; PERMANOVA, *p* < 0.05; Fig. 3A; Table A.5). The diversities of prokaryotic communities of biofilms exposed to the two highest Co concentrations (5 x 10^-7^ and 1 x 10^-6^ M) were significantly different only after 28 d (Unweighted UniFrac; PERMANOVA, *p-* < 0.05; Fig. 3A; Table A.5). Taking temporal dynamics into account, the prokaryotic community composition of biofilms in the control group were different between 1 h and 7, 14, 21 and 28 d (Unweighted UniFrac; PERMANOVA, *p* < 0.05; Fig. 3A; Table A.5). Prokaryotic community composition for biofilms exposed to Co were different between hour 1 and day 1, yet were similar between hour 1 and day 3. As the experiment continued, compositions on day 3 differed from those on day 7, whereby the composition did not change significantly for the remainder of experimentation (day 28) (Unweighted UniFrac; PERMANOVA, *p* < 0.05; Fig. 3A; Table A.5). It is important to note that the diversity of all prokaryotic communities differed significantly from those of the control biofilms after 14 d of exposure (Weighted UniFrac; PERMANOVA, *p* < 0.05; Fig. 3B; Table A.5).

**Fig. 3:**
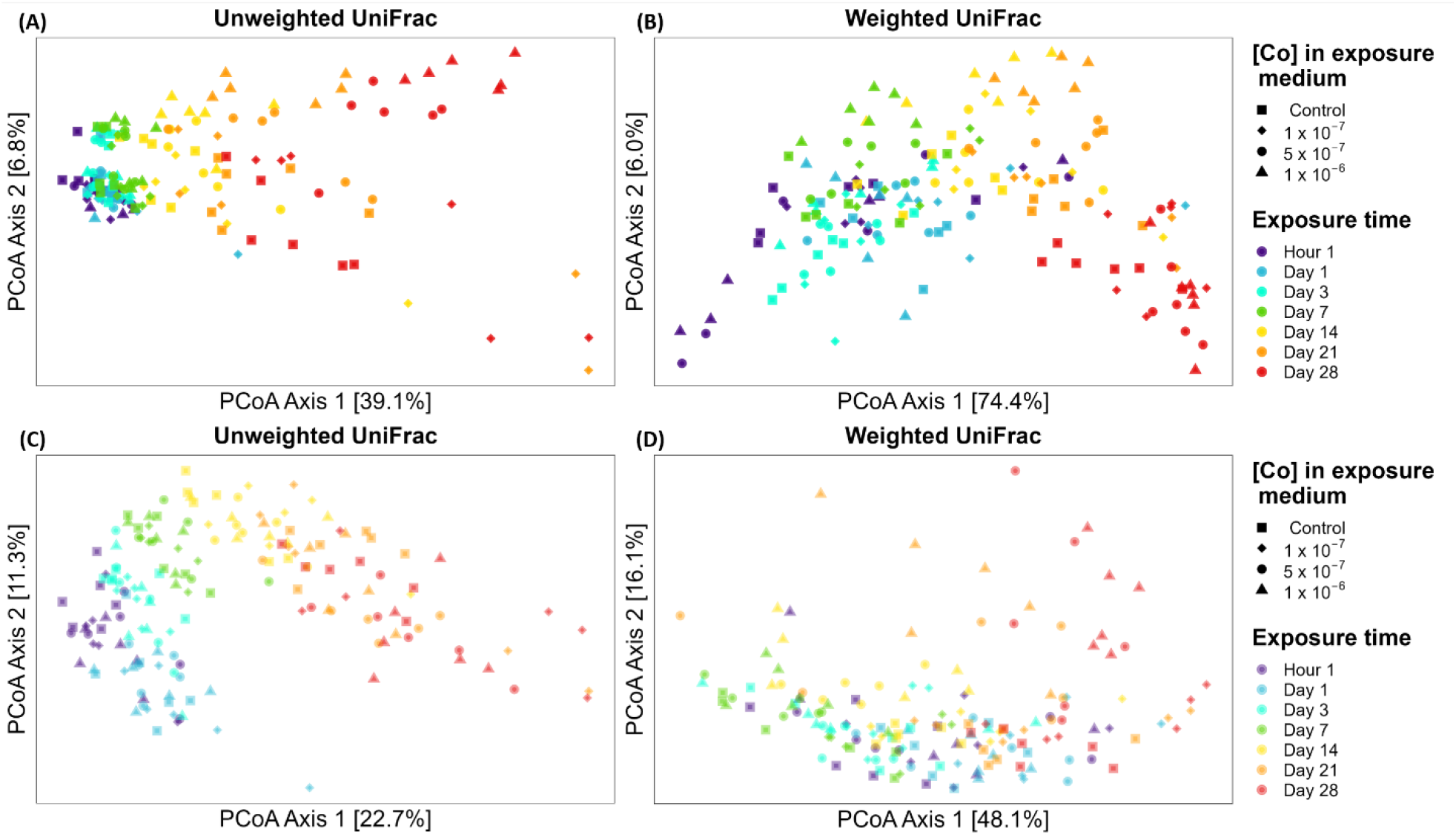
Effects of cobalt concentrations and exposure time on biofilm composition and diversity. (A) PCoA on prokaryotic community (Unweighted UniFrac distance). (B) PCoA on prokaryotic community (Weighted UniFrac distance). (C) PCoA on eukaryotic community (Unweighted UniFrac distance). (D) PCoA on eukaryotic community (Weighted UniFrac distance). (color should be used)

#### 3.3.3. Eukaryotic biofilm community analysis

Analysis of eukaryotic community composition within the V8-V9 region of the 18S rRNA-encoding gene of the 192 biofilm metabolite-extract samples yielded 46,994,861 raw reads, of which 73.4% were retained after sequence assembly, denoising and chimera removal, and 67.8% were retained after a clustering filter (cluster with at least 0.005% of all sequences). Samples displayed 76,931 to 398,084 reads (230,159 ± 49,366). Finally, 902 eukaryotic ASVs were taxonomically assigned.

At the first sampling time (1-h exposure), the richness of control biofilms was 745 ± 15 ASVs (Fig. A.4A) and remained constant throughout the remainder of the experiment. At day 28, Co exposure induced a significant decrease by 14.9%, 10.7% and 17.1% in biofilms exposed to 1 x 10^-7^ M, 5 x 10^-7^ M and 1 x 10^-6^ M, respectively (*post-hoc* Dunn test, *p* < 0.05). The evenness was also similar in control biofilms over time, with a median value of 0.57 (Fig. A.4B). In biofilms exposed to 5 x 10^-7^ M and 1 x 10^-6^ M Co, a significant increase in evenness was observed by as early as day 14. At day 28, Co concentration induced a 15.2% and 17.1% significant increase in evenness, respectively (*post-hoc* Dunn test, *p* < 0.05). The Shannon index was also similar in control biofilms over time, with a median value of 3.80 (Fig. A.4C). In biofilms exposed to 5 x 10^-7^ M and 1 x 10^-6^ M Co, a significant increase in the Shannon index was observed by day 14, and by day 28, Co concentration induced a significant increase to 4.24 and 4.31, respectively (*post-hoc* Dunn test, *p* < 0.05).

Bacillariophyceae (diatoms) represented more than 50% of the relative abundances in all samples (Fig. A.3B). The remaining classes were mostly Clitellata (11%) and Catenulida (11%). Visually, samples from different time points were well dispersed along the first axis of the PCoA, and exposure concentrations were rather separated on the second axis of the PCoA (Fig. 3C and 3D). After 1 h of exposure, the composition and the diversity of eukaryotic communities were not impacted by Co exposure. From day 14 to the end of the experiment, eukaryotic community compositions were significantly different between those of control biofilms and those of biofilms exposed to the two highest concentrations (5 x 10^-7^ and 1 x 10^-6^ M), while those of biofilms exposed to 1 x 10^-7^ M were only different from the control group on day 14 (Unweighted UniFrac; PERMANOVA, *p* < 0.05; Fig. 3C; Table A.6). From day 14 to day 28, eukaryotic community diversity was significantly different between the control biofilm and the biofilm exposed to the highest concentration (Weighted UniFrac; PERMANOVA, *p* < 0.05; Fig. 3D; Table A.6). At days 14 and 28, eukaryotic community diversities were also significantly different in control biofilm and in biofilm exposed to 5 x 10^-7^ M (Weighted UniFrac; PERMANOVA, *p* < 0.05; Fig. 3D; Table A.6). On days 14, 21 and 28, all eukaryotic community compositions were significantly different between each group of exposed biofilms (except between 1 x 10^-7^ and 5 x 10^-7^ M on day 28) (Unweighted UniFrac; PERMANOVA, *p* < 0.05; Fig. 3C; Table A.6). By day 28, an increase as a function of Co concentrations of relative abundance of Chlorophyceae, Eustigmatophyceae and Ulvophyceae was observed as well as a decrease of Bacillariophyceae and Catenulida (*post-hoc* Dunn test, *p* < 0.05; Fig. A.3B). Taking temporal dynamics into account, all eukaryotic community compositions differed significantly from those of the control biofilms after 1 d of exposure (except for those of biofilms exposed to 5 x 10^-7^ and 1 x 10^-6^ M on day 3) (Unweighted UniFrac; PERMANOVA, *p* < 0.05; Fig. 3C; Table A.6). The diversity of all eukaryotic communities differed significantly from those of the control biofilms after 7 d of exposure (except for those of biofilms exposed to 5 x 10^-7^ and 1 x 10^-6^ M on day 7) (Weighted UniFrac; PERMANOVA, *p* < 0.05; Fig. 3D; Table A.6).

#### 3.3.4 Diatom composition and mortality

Microscopy observations were in agreement with genomic analyses and revealed that diatoms dominated the autotrophic community. Diatom densities significantly decreased only at day 28, from 3.67 ± 0.36 to 2.31 ± 0.65 (−37%) and 1.21 ± 0.55 (−67%) x 10^6^ cells·cm^-2^ at the two highest exposure concentrations, respectively (Fig. 2C, Table A.4) (*post-hoc* Dunn test, *p* < 0.05). A significant increase in diatom mortality was observed on day 21 in biofilms exposed to 1 x 10^-6^ M, rising from 2.02 ± 0.04 in the control group to 6.90 ± 0.21% (+242%) (Fig. 2D, Table A.4) (*post-hoc* Dunn test, *p* < 0.05). Although not significant, the higher Co concentration induced greater diatom mortality at each time point than that observed in the control group. The lowest concentration (1 x 10^-7^ M) had no significant effect on these biological parameters (Fig. 2C, 2D and Table A.4). Cobalt exposure led to changes in diatom community composition, with a decrease in *Achnanthidium* F.T. Kützing spp. and *Cocconeis placentula* Ehrenberg var. *lineata* (Ehr.) Van Heurck and an increase in *Eolimna minima* (Grunow) Lange-Bertalot, *Fragilaria capucina* Desmazièrese var. *capucina*, *Fragilaria capucina* Desmazières var. *vaucheriae* (Kützing) Lange-Bertalot, *Fragilaria crotonensis* Kitton, *Gomphonema parvulum* (Kützing) Kützing var. *parvulum* f. *parvulum* and *Mayamaea atomus* var. *permitis* Hustedt Lange-Bertalot (Fig. A5 and Table A.7).

### 3.4. Biofilm meta-metabolomic fingerprints

#### 3.4.1. Untargeted meta-metabolomic features

In total, 5,141 untargeted meta-metabolomic features were observed among all samples while only 192 (3.7%) annotations were attempted using the molecular network approach supported by public spectral database inquiry. However, over 92% of the global meta-metabolome appeared significantly impacted during the 28 d of exposure. Cobalt concentrations had a significant effect on the contents of 2,356 untargeted meta-metabolomic features, exposure time on the contents of 4,703 untargeted meta-metabolomic features and their interaction induced a change in the contents of 1,950 untargeted meta-metabolomic features (two-way ANOVA, *p* <0.05).

A gradual change over time in the meta-metabolomic fingerprints of biofilms with the influence of Co exposure concentrations was observed (Fig. A.6A). Except at day 7, the meta-metabolomic fingerprints of control biofilms at each time point studied were different from those at the first-time point (hour 1) (PERMANOVA, *p* < 0.05; Table A.8). After 1 h of exposure, the meta-metabolomic fingerprints were already different between the control and Co-exposed biofilms (PERMANOVA, *p* < 0.05; Table A.8). On days 1, 3, 7 and 14, the meta-metabolic fingerprints of biofilms showed no significant differences between control and exposed biofilms (PERMANOVA; Table A.8). At day 21, the meta-metabolomic fingerprints were significantly different when comparing control biofilms and biofilms exposed to 5 x 10^-7^ and 1 x 10^-6^ M Co (PERMANOVA, *p* < 0.5; Table A.8). After 28 d of exposure, the meta-metabolomic fingerprints of the control biofilms and biofilms exposed to 1 x 10^-7^ M were significantly different from those of the biofilms exposed to the two highest Co concentrations (5 x 10^-7^ and 1 x 10^-6^ M) (PERMANOVA, *p* < 0.05; Table A.8). Moreover, the meta-metabolomic fingerprints of biofilms exposed to 5 x 10^-7^ M were also significantly different from those of biofilms exposed to 1 x 10^-6^ M (PERMANOVA, *p* < 0.05; Table A.8).

#### 3.4.2. Annotated metabolites in biofilms

Over 97% of the 192 annotated metabolites were significantly impacted during the 28 days of exposure. Cobalt concentrations had a significant effect on the levels of 143 metabolites, while exposure time on the levels of 186 metabolites and their interaction induced a change in the levels of 123 metabolites (two-way ANOVA, *p* <0.05).

Various families of metabolites were observed, such as flavonoids, amino acids, nucleic acids and many lipids or lipid precursors, including lyso-diacylglyceryltrimethylhomoserines (LDGTS), lysophosphatidylcholines (LPC), phosphatidylethanolamines (PE) and steroids (Fig. A.7 and Table A.9). Globally, considering the entire meta-metabolome, similar trends were observed for the 3.7% of untargeted meta-metabolomic features that have been annotated (Table A.9). The application of similar data processing better evidenced the influence of Co concentration and exposure time using the whole meta-metabolome data set than the 192 annotated metabolites (Fig. A.6B).

#### 3.4.3 Determination of CRIDeR/CRIDaR based on untargeted meta-metabolomic features

Dose-response models were built based on the peak intensity of the untargeted meta-metabolomic features as a function of added Co concentrations in exposure media for each time point. They were then classified into four different categories as a function of their respective curve trends. In total, the dose-response curves of 285 features exhibited a bell-shaped trend, 118 a U-shaped trend, 1051 an increasing trend and 783 a decreasing trend (Fig. 4A and Table A.10). Using ECDF curves to aggregate (Fig. 4B) the BMDs retrieved from those dose-response curves, CRIDeR initiation thresholds were determined from the best fit model at each time point (Table A.11). These thresholds correspond to the hazardous concentrations at which 5% of untargeted meta-metabolomic features with a biphasic dose-response trend will potentially be affected on the basis of their BMDs (Fig. 4C and A.8). The Co concentrations initiating CRIDeRs were found to be comparable at each exposure time: 1.55 x 10^-10^, 8.65 x 10^-11^, 1.14 x 10^-11^, 2.26 x 10^-10^, 1.69 x 10^-10^, 2.48 x 10^-10^ and 9.23 x 10^-11^ M of Co^2+^ added in exposure medium, respectively (Fig. 4, A.8 and Table 1). The CRIDaR initiation thresholds were also determined from the best fit model at each time point (Table A.11) and correspond to the hazardous concentrations at which 5% of untargeted meta-metabolomic features with a monotonic dose-response trend will potentially be affected on the basis of their BMDs (Fig. 4C and A.8). The Co concentrations initiating CRIDaRs increased gradually over time; starting at 1.93 x 10^-9^ by hour 1, increasing to 4. x 10^-9^ by day 3 and 6.40 x 10^-9^ by day 14, ending at 2.00 x 10^-8^ and 1.22 x 10^-8^ M by days 21 and 28, respectively (Fig. 4, A.8 and Table 1).

**Fig. 4:**
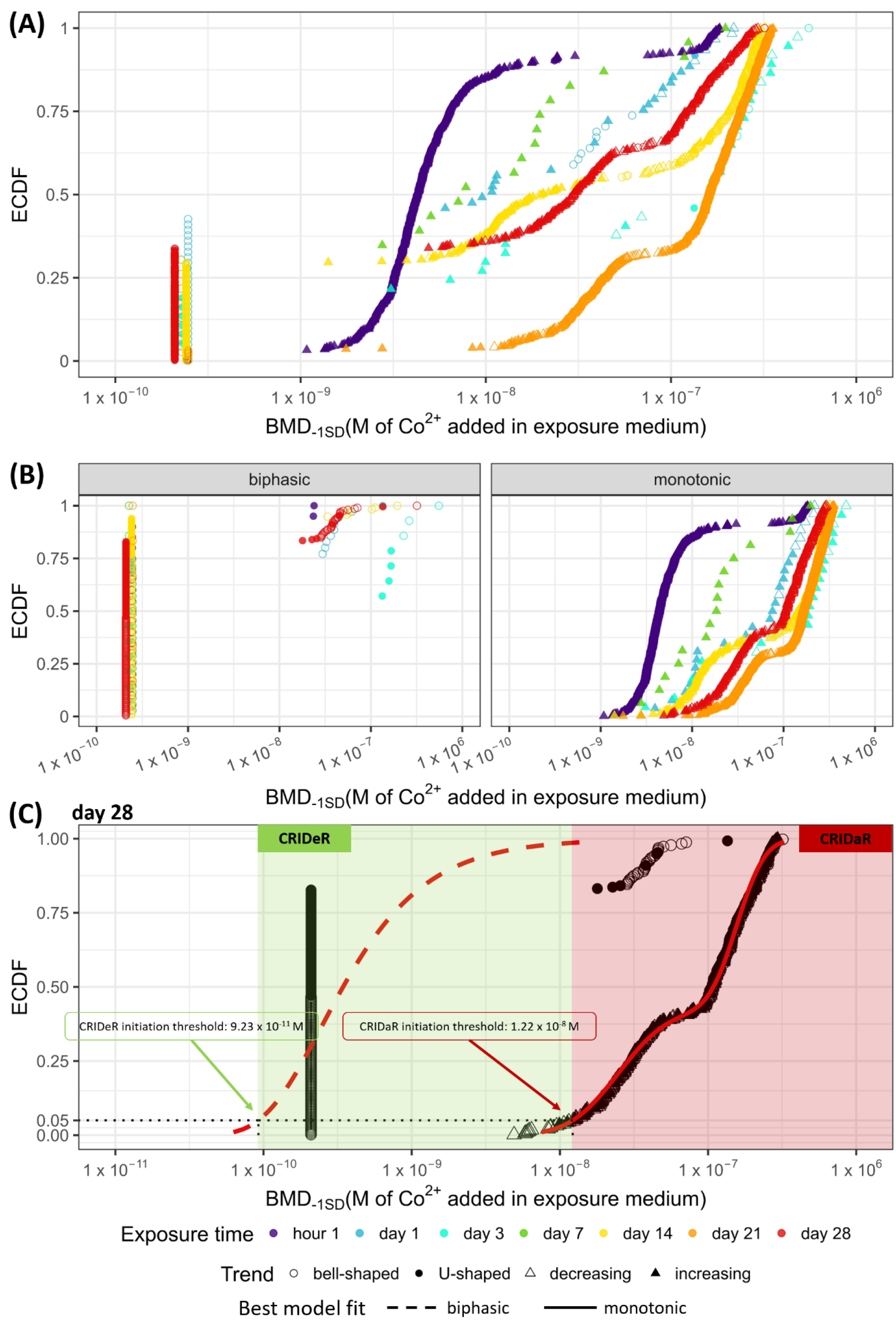
Biofilm meta-metabolomic responses to Co concentrations. (A) Distribution of BMD_-1SD_ (M of Co^2+^ added in exposure medium) as an ECDF at each sampling time step. (B) Distribution of BMD_-1SD_ (M of Co^2+^ added in exposure medium) as an ECDF split by trend of dose-response curve; biphasic (bell- and U-shaped) or monotonic (decreasing and increasing). (C) Determination of CRIDeR and CRIDaR initiation thresholds at day 28 from the best fit model for ECDFs of untargeted meta-metabolomic features with biphasic (bell- or U-shaped) and monotonic (decreasing or increasing) trends, respectively. CRIDeR and CRIDaR initiation thresholds are the hazardous concentration at which 5% of untargeted meta-metabolomic features will potentially be affected on the basis of their BMDs. Determination of CRIDeR and CRIDaR initiation thresholds for the other sampling time are presented in Fig. A.8 and Table 1. (color should be used)

**Table 1:**
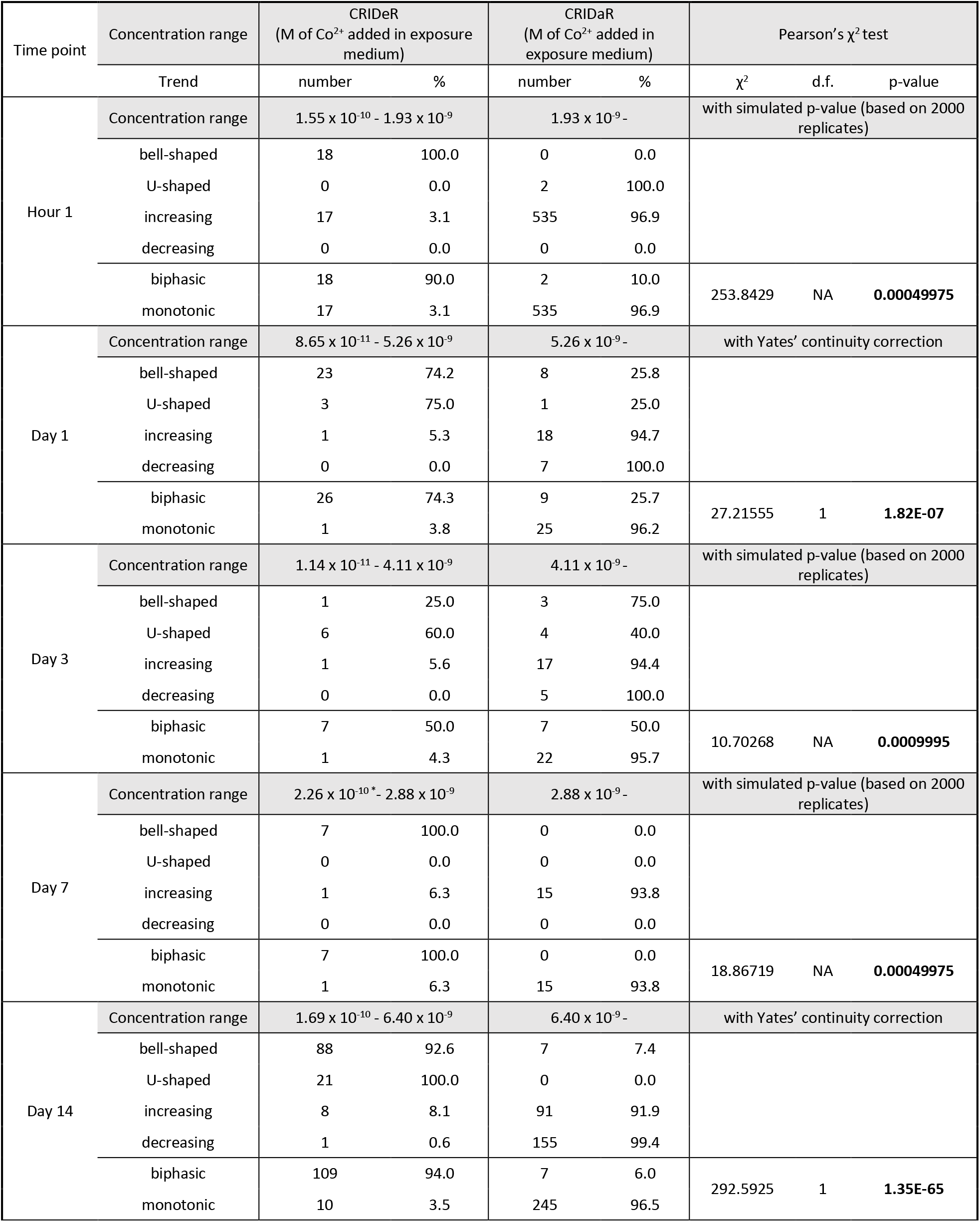
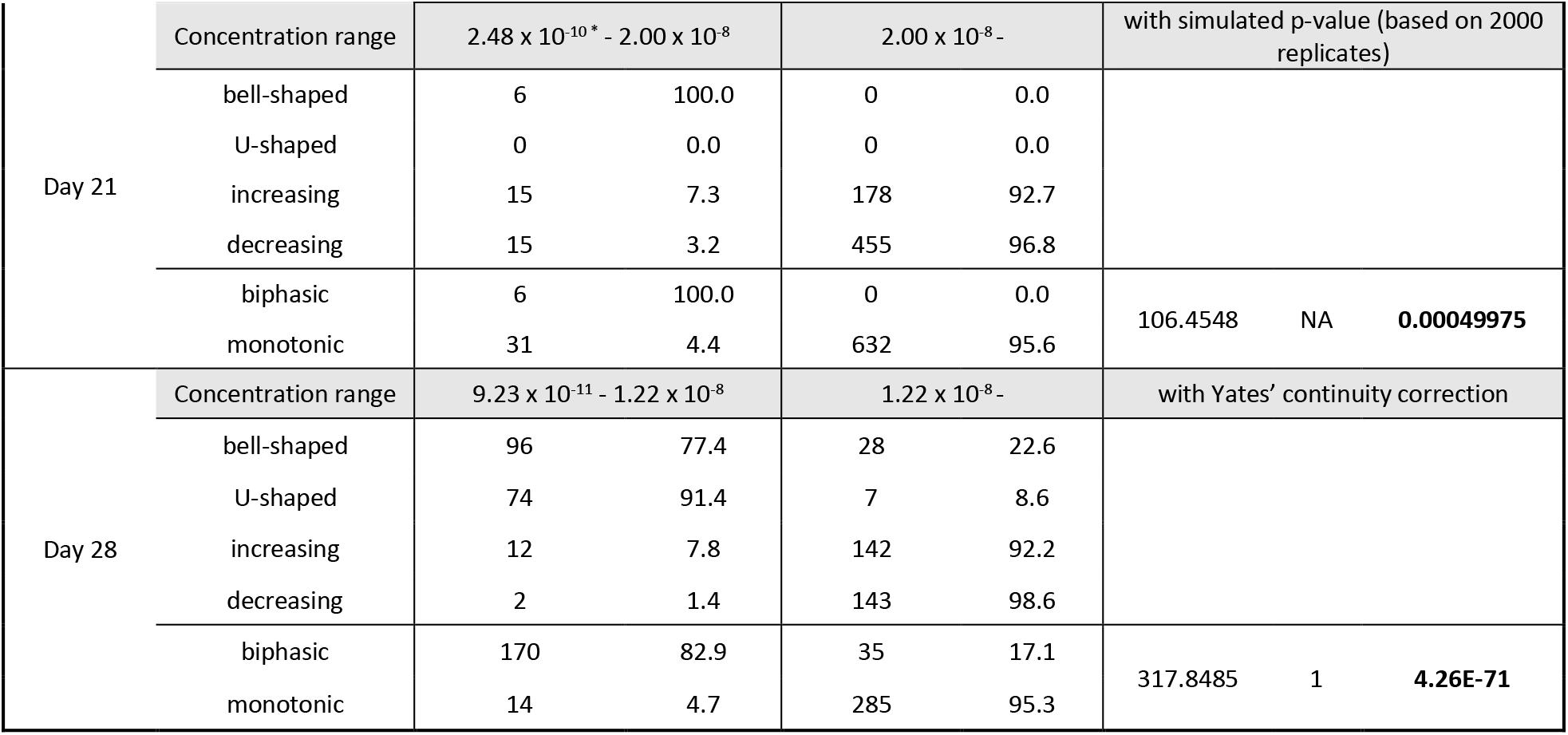
Calculated CRIDeR and CRIDaR at each time point. * CRIDeR threshold determined directly with the BMD of the first untargeted meta-metabolomic feature impacted because all distributions failed to fit.

After 1 h of exposure, 90% of untargeted meta-metabolomic features with biphasic dose-response trends were present in the CRIDeR (Table 1). That percentage decreased to 74.3% and 50.0% at days 1 and 3 (Table 1). By day 7, 100% of the untargeted meta-metabolomic features with biphasic dose-response trends were present in CRIDeR, whereas 94.0%, 100% and 82.9% were present at 14, 21 and 28 d, respectively (Table 1). At each time point, 96.9%, 96.2%, 95.7%, 93.8%, 96.1%, 95.3% and 95.3%, respectively, of untargeted meta-metabolomic features with monotonic trends were present in CRIDaR (Table 1). The proportions of untargeted meta-metabolomic features with biphasic dose-response trends (bell-shaped and U-shaped) were significantly greater in CRIDeR than in CRIDaR (Pearson’s χ^2^ test, *p* < 0.001; Table 1). Conversely, the proportions of untargeted meta-metabolomic features with monotonic dose-response trends (increasing and decreasing) were significantly greater in CRIDaR than in CRIDeR (Pearson’s χ^2^ test, *p* < 0.001; Table 1).

#### 3.4.4. Determination of TRIDeR/TRIDaR based on untargeted meta-metabolomic features

Time-dose models were constructed based on the intensity of the metabolomic feature peaks as a function of time for each exposure with Co^2+^ added to the exposure medium. The curves were further classified into four different categories as a function of their respective time-response model curve trend. In total, 1,784 exhibited a bell-shaped trend, 235 a U-shaped trend, 5,443 an increasing trend and 177 a decreasing trend (Fig. 5 and Table A.12). Using ECDF curves to aggregate (Fig. 5B) the BMTs retrieved for those dose-response curves, TRIDeR initiation exposure times were determined from the best fit model at each time point (Table A.13), and correspond to the hazardous times where 5% of untargeted meta-metabolomic features with a biphasic dose-response trend will potentially be affected on the basis of their BMDs (Fig. 5C and A.9). TRIDeR initiation times were found to be mostly instantaneous at each exposure concentration with a value of 17, 16 and 17 s, for the exposure concentrations of 1 x 10^-7^, 5 x 10^-7^ and 1 x 10^-6^ M Co, respectively. Over the first few moments of exposure (36 s), 23.5%, 28.6% and 24.9% of untargeted meta-metabolomic features with a bell-shaped time-response trend were impacted in biofilms exposed to 1 x 10^-7^, 5 x 10^-7^ and 1 x 10^-6^ M, respectively (Fig. 5A, 5B and Table 2). The first untargeted meta-metabolomic feature in the control group was affected after 64 min (Fig. 5 and Table 2).

**Fig. 5:**
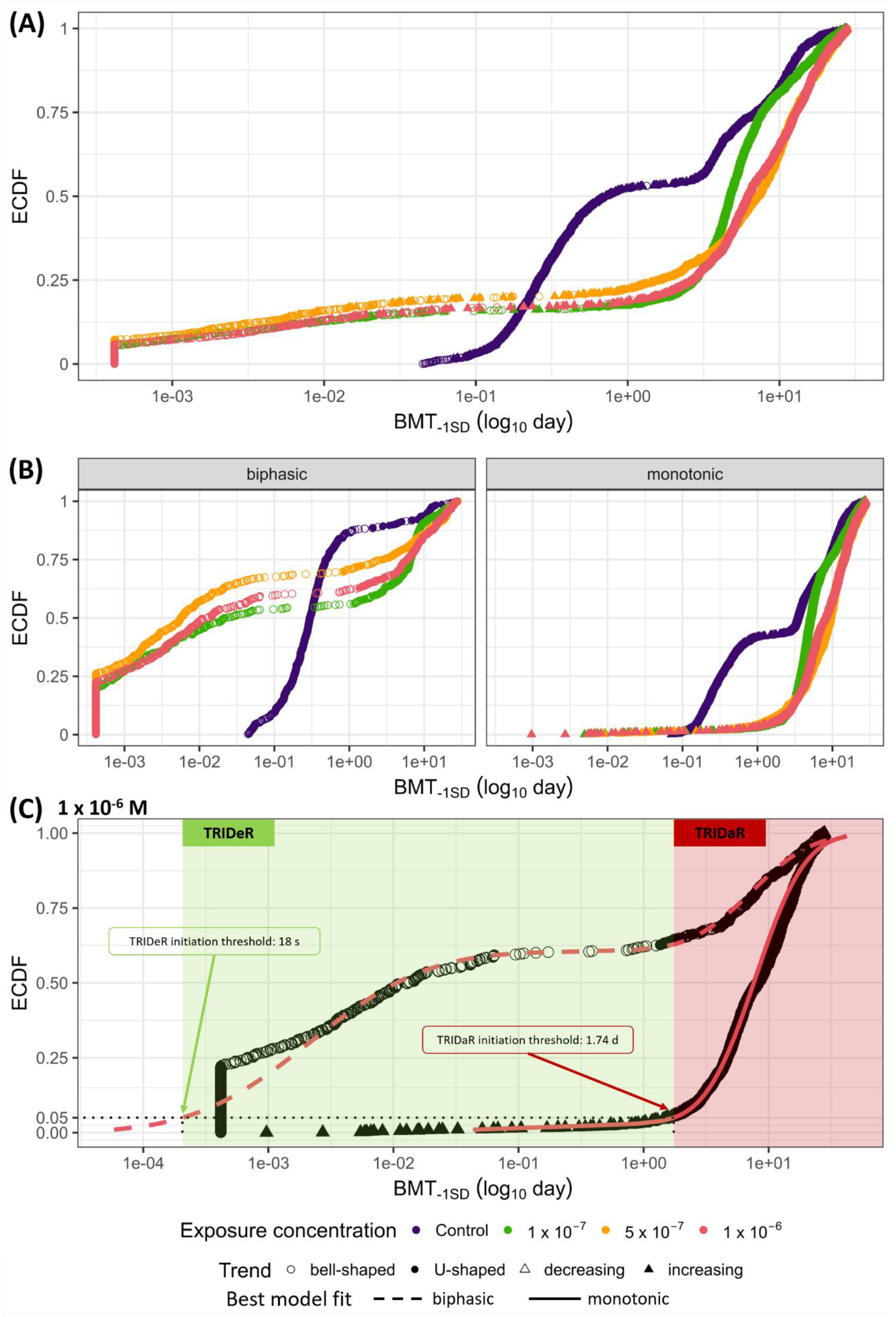
Biofilm meta-metabolomic responses to Co exposure times. (A) Distribution of BMT_-1SD_ (in days; log_10_ transformed) as an ECDF for each exposure concentration. (B) Distribution of BMT_-1SD_ (in days; log_10_ transformed) as an ECDF split by trend of dose-response curve; biphasic (bell- and U-shaped) or monotonic (decreasing and increasing). (C) Determination of TRIDeR and TRIDaR initiation thresholds for 1 x 10^-5^ M exposure concentration from the best fit model for ECDFs of untargeted meta-metabolomic features with biphasic (bell- or U-shaped) and monotonic (decreasing or increasing) trends, respectively. TRIDeR and TRIDaR initiation thresholds are the hazardous exposure time at which 5% of untargeted meta-metabolomic features will potentially be affected on the basis of their BMTs. Determination of TRIDeR and TRIDaR initiation thresholds for the other sampling times are presented in Fig. A.9 and Table 2. (color should be used)

**Table 2:**
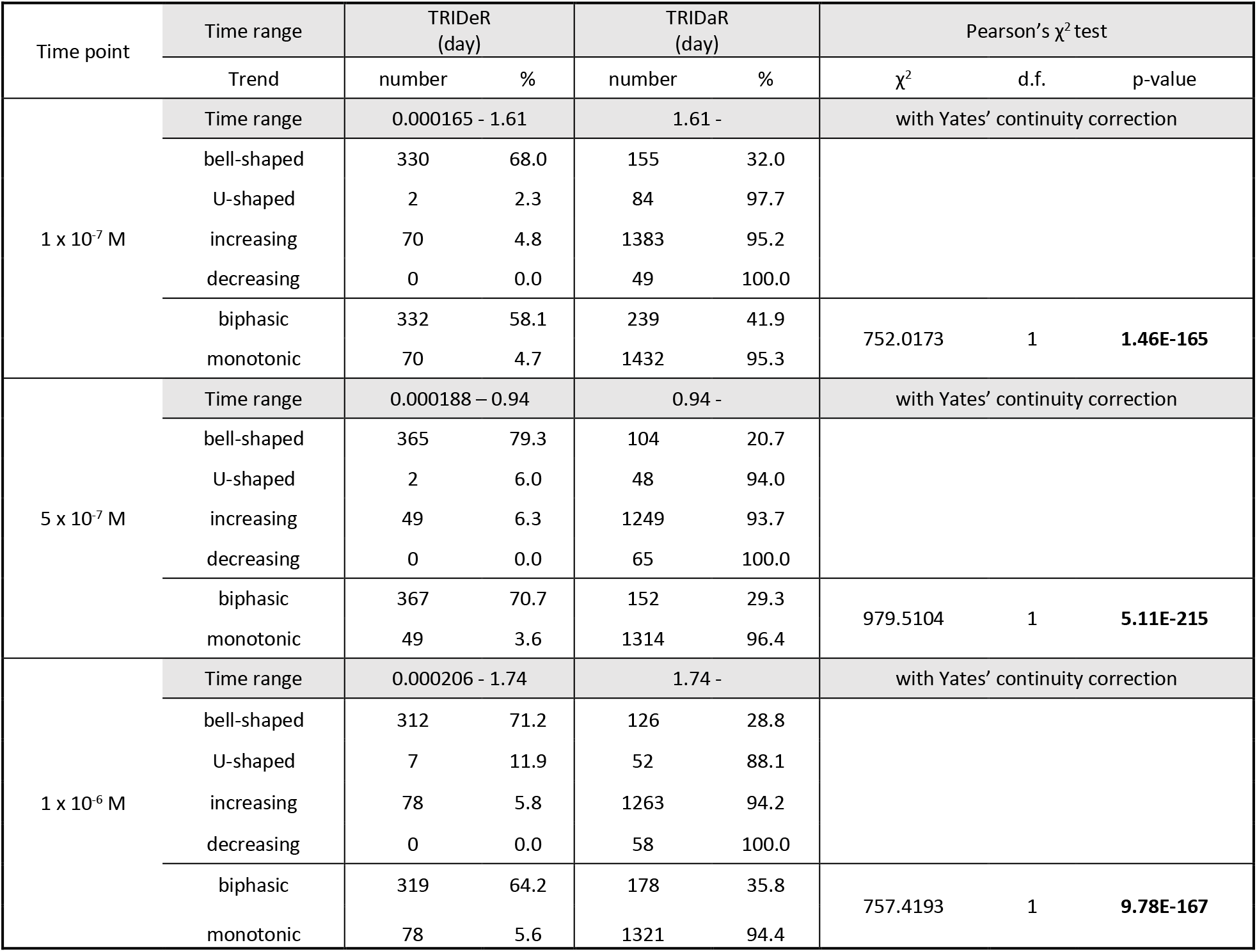
Calculated TRIDeR and TRIDaR for each Co exposure concentration group.

More than half (58.1%) of untargeted meta-metabolomic features with a biphasic trend and 4.7% with a monotonic trend were impacted within the TRIDeR for biofilms exposed to 1 x 10^-7^ M (Table 2). Similar results were observed for biofilms exposed to 5 x 10^-7^ and 1 x 10^-6^ M, with 72.3% and 64.2% of untargeted meta-metabolomic features with a biphasic trend, respectively, and 6.0% and 5.6% with a monotonic trend also impacted within the TRIDeR (Table 2). The TRIDaR initiation times appeared after one day at each exposure concentration: 1.61 d, 1.44 d and 1.74 d, respectively (Fig. 5A and Table 2). The ECDF profiles were similar between exposure concentrations (1 x 10^-7^, 5 x 10^-7^ and 1 x 10^-6^ M groups) and different from those of the control group (Fig. 5A and 5B). The proportions of untargeted meta-metabolomic features with biphasic dose-response trends were significantly greater in TRIDeR than in TRIDaR (Pearson’s test, *p* < 0.001; Table 2). Conversely, the proportions of untargeted meta-metabolomic features with monotonic dose-response trends were significantly greater in TRIDaR than in TRIDeR (Pearson’s χ^2^ test, *p* < 0.001; Table 2).

### 3.6. Omic responsiveness kinetics

The global metabolite content changes (expressed as a % of the total composition change trajectories over 28 days) observed in the meta-metabolome occurred more rapidly than those observed in the prokaryotic and eukaryotic community compositions, regardless of Co exposure concentrations (Fig. 6A-D). The early change (1 h) in the meta-metabolome due to Co exposure was 20.8, 21.8 and 23.7% (of the total change observed during the 28 days) compared with the control biofilms exposed to 1 x 10^-7^, 5 x 10^-7^ and 1 x 10^-6^ M, respectively. At the same time point, the kinetics of change in the prokaryotic community were 11.1, 11.6 and 14.3%, respectively, and 12.6, 12.5 and 12.4%, respectively, in the eukaryotic community. In the control group, the kinetics of composition change were faster in the eukaryotic community than in the prokaryotic community (Fig. 6A). Similar rates of change were observed in biofilm communities exposed to 1 x 10^-7^ M, where the kinetics of change were also faster in the 18S-derived community than in the 16S-derived community (Fig. 6B). However, for biofilms exposed to 1 x 10^-6^ M, the prokaryotic community was impacted more rapidly than the eukaryotic community (Fig. 6D). This transition in change kinetics was gradual, since for biofilms exposed to 5 x 10^-7^ M, the change kinetics of the two communities were similar up to day 3, with less difference thereafter compared to the control and 1 x 10^-7^ M Co exposure groups (Fig. 6C).

**Fig. 6:**
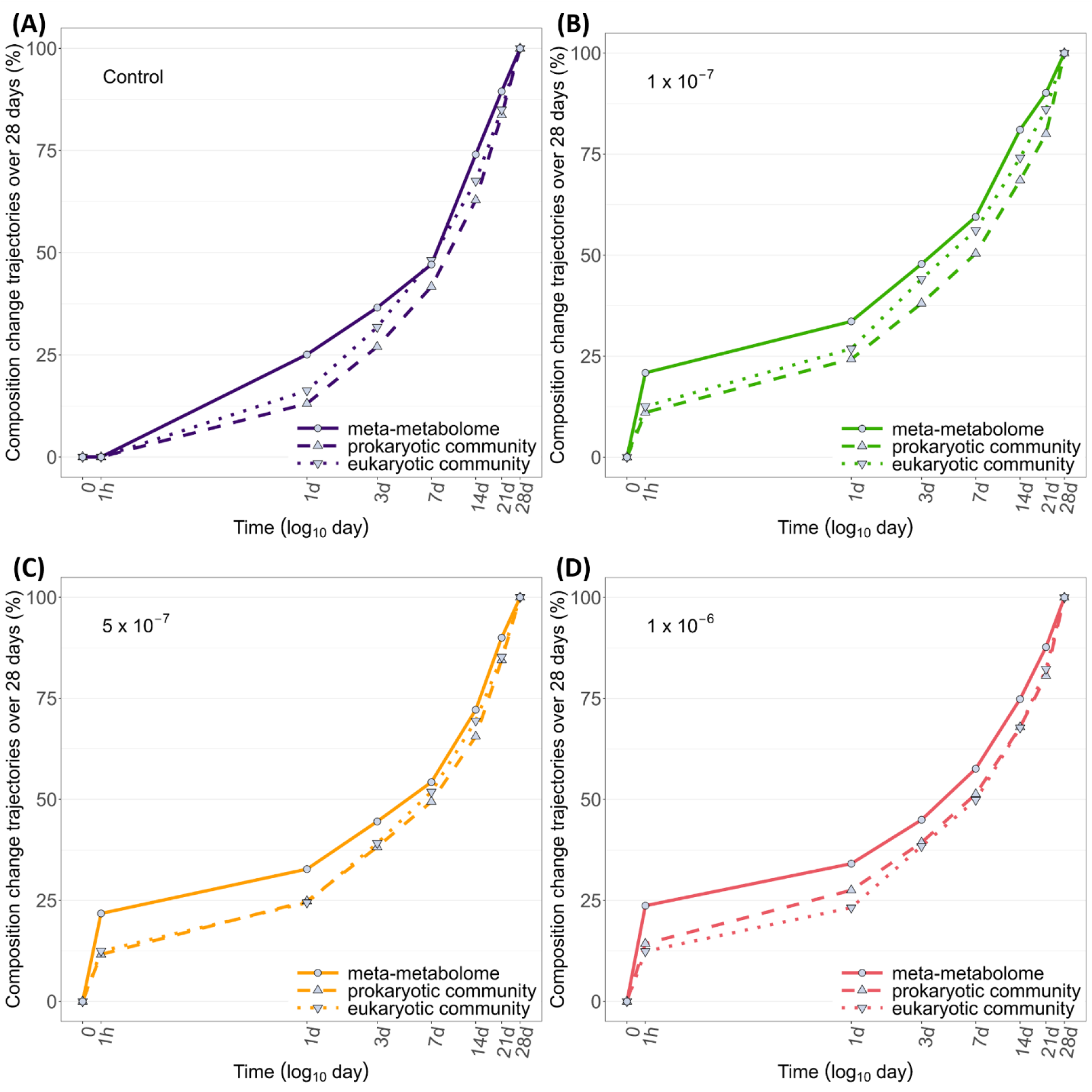
Omics kinetics based on dissimilarities between biofilm meta-metabolomes (Fig. A.10A), prokaryotic (Fig. A.10B) and eukaryotic (Fig. A.10C) communities as a function of Co concentration and exposure time. Composition change trajectories in meta-metabolome (solid line), prokaryotic community (dashed line) and eukaryotic community (dotted line) expressed as percentages of the total trajectory achieved at hour 1, days 1, 3, 7, 14, 21 and 28; (A) for biofilms of control group, (B) for biofilms exposed to 1 x 10^-7^ M Co, (C) for biofilms exposed to 5 x 10^-7^ M Co and (D) for biofilms exposed to 1 x 10^-6^ M Co. (color should be used)

## 4. Discussion

### 4.1. Concentrations and exposure time to Co induce change in biofilm structure

Cobalt accumulation in biofilms was quantifiable as early as 1 h after exposure (Fig. 1, Fig. A.1 and Table A.2). As with other metals such as Cu, Zn or Ni (Meylan et al., 2003; Laderriere et al., 2020), a correlation between free Co^2+^ ion concentrations and intracellular Co was observed (Fig. A.1). This bioaccumulation reflects the bioavailability of Co to biofilms, and therefore its potential toxic effects (McGeer et al., 2004).

Control biofilms had specific temporal dynamics over time. An initial increase in biomass was observed during the first week, followed by a gradual decrease until day 21, and an increase again on day 28 (Fig. 2A). The increase in chlorophyll *a* content over the 28-d experiment appeared to be closely linked to that of diatom density (Fig. 2B) consistent with their dominance in relative abundance of the eukaryotic community (Fig. A.3B). A linear regression analysis between the total diatom abundance and chlorophyll *a* content showed a significant positive correlation relationship (Patil and Anil, 2005). An overall change in diversity compared with the initial biofilms was observed over time (without taking into account the exposure concentration factor) for the prokaryotic community after 7 d and after 1 d for the eukaryotic community. These changes therefore appear to be initiated naturally by environmental factors. In fact, the experiment took place in autumn, and water temperatures and sunshine decreased progressively throughout the month of exposure (Table A.1). Indeed, after a fall biomass peak, a decrease in specific richness can be observed in river biofilms (Lyautey et al., 2005; Morin et al., 2008a). These observations are also supported by the progressive decrease of alpha diversity indices within the control group, *i.e.* in richness (Fig. A.2A), evenness (Fig. A.2B) and Shannon (Fig. A.2C) for prokaryotes and in richness (Fig. A.4A) for eukaryotes.

The structure of biofilms chronically exposed to Co was both rapidly and lastingly affected. With regard to traditional biological monitoring parameters, the biomass of biofilms exposed to 1 x 10^-6^ M Co began to decrease significantly compared with that of the control group after two weeks of exposure, as did chlorophyll *a* content (Fig. 2A, 2B and Table A.4). These results were in line with those observed following exposures to other metals (Massieux et al., 2004). Surprisingly, the diversity of biofilm communities was rapidly affected by exposure to Co. As early as 1 h after exposure, the communities of prokaryotes exposed to 5 x 10^-7^ and 1 x 10^-6^ Co were lower than those of the control group. However, after 1 d of exposure, these differences were no longer detected. As observed for traditional parameters (biomass and chlorophyll *a* content), prokaryotic and eukaryotic communities were significantly and durably impacted by Co after 14 days of exposure (Fig. 3, A.3, A.5 and Tables A.5, A.6 and A.7). This is in line with the effects of metals on microbial community structures in biofilms. After 21 days of exposure to Ag, structures of bacterial and algal communities were both impacted by exposure conditions (Gil-Allué et al., 2018).

In control biofilms, the eukaryotic community changes more rapidly than the prokaryotic community (Fig. 6A). This is also observed in biofilms exposed to 1 x 10^-7^ M Co (Fig. 6B). On the other hand, Co seems to influence these dynamics of community changes and succession. In biofilms exposed to 5 x 10^-7^ M Co, both communities had similar change kinetics on microbial diversity for up to 3 days of exposure (Fig. 6C). This effect was even greater in biofilms exposed to 1 x 10^-6^ M Co, where the change kinetics proved to be faster for the prokaryotic than the eukaryotic community in terms of taxa occurrence (Fig. 6D). Cobalt would therefore have a faster adverse effect on prokaryotic communities, very likely by inducing the rapid disappearance of the most sensitive taxa.

### 4.2. Changes in the meta-metabolome are faster than those observed in prokaryotic and eukaryotic community compositions

The comparison of variations at different levels of biological response through different omic approaches (Fig. 6) underlined the value of performing first metabolite extraction prior to DNA extraction on the same sample (Duperron et al., 2023) as applied here for ecotoxicological investigations. Indeed, the three parameters studied (metabolites, 16S and 18S ASVs) came from the same samples, and provided a good representation of the response of microbial communities to contamination on individualized biofilm samples. As emphasized recently, such combination of omics approaches is a new direction to take in environmental risk assessment, and particularly in microbial ecotoxicology (Hellal et al., 2023).

As for the changes observed in biofilm prokaryotic and eukaryotic community structures, the meta-metabolome was rapidly impacted (after 1 h) by Co (Table A.8). Then, on days 1, 3 and 7, the meta-metabolomic fingerprint no longer differed between exposed biofilms and those in the control group. This was followed by a more lasting impact from 14 days of exposure to the end of the experiment (Fig. A.6). The metabolomic fingerprints of two freshwater microalgae exposed to Cd and Pb were also different from the control group after 2 weeks (Nanda et al., 2021). In a soil, in which the prokaryote community was exposed for 40 days to trinitrotoluene, cyclotrimethylene trinitramine and cyclotetramethylene tetranitramine (alone or in mixtures), the meta-metabolomic fingerprints were also different between exposure groups (Yang et al., 2021), underlining the fact of the lasting effects of stress on the chemical signature of microbial communities.

Cobalt concentrations, exposure time and natural community succession led to progressive changes in the prokaryotic and eukaryotic communities and meta-metabolome of the biofilms. However, on the basis of the percentages of overall change over 28 days deduced from MOTA, the change in the meta-metabolome appeared the most rapid (Fig. 6). Indeed, when comparing the kinetics of change in both control and exposed biofilms, the meta-metabolome changed more rapidly at each time point than the microbial community structure. Within 1 h of exposure, 20.9, 21.8 and 23.7% of the total meta-metabolome variation had already occurred for biofilms exposed to 1 x 10^-7^, 5 x 10^-7^ and 1 x 10^-6^ M Co respectively, compared to the variation of 11.1, 11.6 and 14.3% for the prokaryotic communities and of 12.6, 12.5 and 12.4% for the eukaryotic communities (Fig. 6). River biofilms exposed to erythromycin and Ag nanoparticles had also showed a change in their meta-metabolome, while prokaryotic community diversity appeared unaffected after 6 and 12 weeks of exposure, respectively (Pu et al., 2021; Yang et al., 2023).

These observations support the view that the (meta-)metabolomic fingerprint constitutes an instantaneous illustration of the molecular response of organisms in their environment (see section 4.4). Also, this metabolite fingerprint, which reflects the physiological status of biological systems, appears here even more sensitive than the specific response at the level of the composition of microbial community (see section 4.3).

### 4.3. CRIDeR and CRIDaR remain relevant even in a context of changing and **“tolerant” communities**

The interpretation of trends of dose-response models of untargeted meta-metabolomic features and the identification of CRIDeR and CRIDaR may provide fundamental information on the level of biofilm responses to a stress (Colas et al., 2023). One can expected that these CRIDeR and CRIDaR would be impacted by community changes over time and the appearance of tolerant and resistant species, leading to an increase in their tolerance threshold.

However, the determination of the different CRIDeRs observed at the different time point suggested that defense responses were initiated as soon as the most sensitive microorganisms of the biofilms are first disturbed by an addition of Co in the exposure media. Indeed, the CRIDeRs calculated throughout the experiment had similar initiation threshold values of above 1.41 ± 0.77 x 10^-10^ M of Co^2+^ added in the exposure medium (Table 1). At Co^2+^ concentrations above background concentrations (1.36 ± 0.85 x 10^-9^ M Co^2+^), the metabolic processes of biofilms reacted and appeared to set up molecular defense responses in order to manage Co uptake (Fig. 1, A.1 and Table A.2). These defense responses took place at lower concentrations than those previously observed in static microcosms (Colas et al., 2023). It is well known that contaminants can impact the (meta-)metabolome of biological systems. At 4.3 x 10^-10^ M of diuron, biofilm meta-metabolomic fingerprint also began to be impacted as determined via BMD calculation on DRomics (Creusot et al., 2022).

The CRIDeRs of the microbial communities were stable despite a change in taxonomic structure induced by the Co exposure time, which is most likely due to the influence of external environmental factors in addition to the effects induced by Co concentrations (Fig. 3, 4, A.3, A.5, Tables 1, A.5, A.6 and A.7). After 28 days of exposure to Co, the biofilm communities have probably been able to acclimatize to this disturbance, and be characterized by the presence of so-called tolerant species. An increase in the relative abundance of previously reported as potential metal-tolerant diatom species (Fig. A.5 and Table A.7), such as *Eolimna minima*, *Fragilaria capucina* var. *capucina*, *Fragilaria capucina* var. *vaucheriae*, *Fragilaria crotonensis*, *Gomphonema parvulum* var. *parvulum* f. *parvulum* or *Mayamaea atomus* var. *permitis* was observed on day 28 (Duong et al., 2008; Tlili et al., 2011; Morin et al., 2012). Also, a decrease in microbial species known to be metal-sensitive was also observed with increasing Co concentrations, such as *Cocconeis placentula* var. *lineata*. In a previous study (Colas et al., 2023), Co induced a change in the composition of photosynthetic pigments, with an increase in the proportion of chlorophyll *b* compared with chlorophylls *a* and *c*_1_+*c*_2_, possibly indicating the favored presence of green algae. On day 28, the greater presence of Chlorophyceae and Ulvophyceae in eukaryote communities in biofilms exposed to the two highest concentrations would thus confirm the metal resistance of certain green algae (Ivorra et al., 2000; Corcoll et al., 2011).

Despite microbial community changes occurring during the 28 days exposure time series, such as an increase in tolerant species and a decrease in sensitive species, the CRIDeR observed after 1 h or after 28 d present both very similar Co initiation-threshold concentrations at above 1.55 and 0.92 x 10^-10^ M of Co^2+^ added in the exposure medium, respectively (Table 1). Trend analyses of the dose-response curves of untargeted meta-metabolomic features therefore seems relevant regardless of exposure time, even in a context of change and succession of prokaryotic and eukaryotic communities. Despite changes in communities (Fig. 3, A.2, A.3, A.4 and A.5), the presence of metal-tolerant species and the disappearance of metal-sensitive species (Table A.7), the concentrations at which defensive responses are initiated (Table 1) remain similar over time. Interestingly, defense mechanisms such as superoxide dismutase activity were observed continuously (from 6 h to 21 d) during exposure to ZnO nanoparticles in river biofilms (Hou et al., 2016).

As Co concentrations increased, a CRIDaR was also characterized at each time point with increasingly higher Co initiation-thresholds (Table 1). While biofilms maintained similar CRIDeR initiation thresholds over time (Table 1), the CRIDaR thresholds increased. This could indicate that the tolerance of microbial communities to Co is manifested by greater resistance to concentrations. Molecular damage would thus occur at higher concentrations after 28 days of exposure than after 1 hour or 1 day (Fig A.8A, A.8G and Table 1). At the same time, defensive responses are still in place from the smallest increases in Co concentration in the environment (Fig A.8A and A.8G).

Note that the modeling of dose-response curves, from which BMDs were calculated, ECDFs constructed and CRIDeRs and CRIDaRs determined, was relying on the number of Co concentrations that were tested and integrated within the model. This number of exposure concentrations could represent a limiting factor in this approach, as the observation of biphasic trends in biomarker responses in ecotoxicology was restricted by the number of concentrations tested in the studies (Colas and Le Faucheur, 2023).

### 4.4. TRIDeR and TRIDaR highlight rapid responses to Co exposition

As expected, the calculation of TRIDeR and TRIDaR following the same approach as previously developed for CRIDeR/CRIDaR and considering the exposure time factor instead of concentrations, is feasible and provides relevant findings. Indeed, whatever the Co concentration tested, exposed biofilms exhibited a TRIDeR initiation threshold of above 17 s (Fig. 5 and Table 2). However, bioaccumulation data (Fig. A.1 and Table A.2) suggested that biofilms began to bioaccumulate Co as early as one hour after exposure. The quick-acting effect of TRIDeR initiation was observed here after only seconds, as expected with the almost immediate metal internalization and potential cellular impact. In fact, the untargeted meta-metabolomic features appear almost instantaneously in each of the concentrations tested, suggesting the occurrence of immediate metabolic effects resultant of intracellular Co increase. Indeed, after 1 min of exposure, 28.0, 31.1 and 28.1% of the whole metabolite feature set were altered in comparison to control (all with a bell-shaped dose-response trend), after exposure to 1 x 10^-7^, 5 x 10^-7^ or 1 x 10^-6^ M Co, respectively (Fig. 5 and Table 2). After 1 h of exposure, these levels of impacted untargeted meta-metabolomic features rose by 60.9, 72.5 and 63.0%, respectively (Fig. 5 and Table 2). In contrast, the first untargeted meta-metabolomic features were impacted only after 1 h in control biofilms (Fig. 5 and Table 2). This shows that, at the metabolome level, biofilms are extremely sensitive to perturbations in the aquatic environment, given their continuous, equilibrium interaction with it. Interestingly, similar rapid responses in the order of tens of seconds were also observed in biofilms exposed to diuron (Creusot et al., 2022). Another study also showed the almost instantaneous (< 1 min) effects of this contaminant on river biofilms and on their photosynthetic activity (Morin et al., 2018), demonstrating the immediate responsiveness of biofilms to such chemical disturbances in its environment. Microalgae also responded rapidly to metal exposure, producing phytochelatins after 15 min of exposure to Cu, Cd and Pb (Morelli and Scarano, 2001, 2004). In river biofilm exposed to ZnO nanoparticles, defense responses were initiated after 6 h of exposure with an increase in the activity of catalase and superoxide dismutase as a function of contaminant concentration (Hou et al., 2016).

The ECDFs of the untargeted meta-metabolomic features of exposed biofilms showed similar temporal profiles at each Co concentration, being largely distinguishable from the slow variations observed in control biofilms (Fig. 5). The TRIDaR initiation thresholds were similar for biofilms exposed to 1 x 10^-7^, 5 x 10^-7^ and 10^-6^ M Co with an exposure time of 1.4 ± 0.4 d. According to the literature, rapid damage effects were also observed on wastewater microbial communities after 5 h of exposure to graphene oxide, with a significant decrease in metabolic activity that also appears to be proportional to increasing contaminant concentrations (Ahmed and Rodrigues, 2013). Taken together, one can deduce that the microbial biofilms response to acute toxicant exposure comprises both defense and damage mechanisms that are mobilized and inducible within tens of seconds and several hours, respectively.

## 5. Conclusion

The use of the whole meta-metabolomic fingerprint of biofilms appears to be a promising biological compartment to be studied for environmental risk assessment, providing additional and complementary information to traditional monitoring parameters and the information deduced solely from few annotated or targeted metabolites.

CRIDeR and CRIDaR characterizations appear consistent even in the long time-series situation where community dynamics are influenced by both the Co concentrations and the exposure time. In particular, it was observed that potentially tolerant and resistant communities present defense responses similar to those of newly exposed biofilms at similar Co concentrations. Interestingly, these microbial communities, which have probably become tolerant following continuous and long-term exposure, are more resistant to the harmful effects caused by Co.

The TRIDeR and TRIDaR determinations reinforce the fact that the microbial biofilm meta-metabolome is a biological component in close and continuous interaction with the external environment, making it possible to appreciate the rapid rate at which Co affects riparian microbial communities. This study also highlighted the complementarity of multi-omics approaches to better understand the factors driving the responses of a community to stress. Above all, these results confirm the usefulness of analyzing all untargeted metabolomic features with a holistic approach without being restricted by their annotation (and putative functions retrieved from previous observation), proposing CRIDeR/CRIDaR and TRIDeR/TRIDaR as additional new tools for ecotoxicology and microbial ecology studies.

## Author contributions

**Simon Colas**: Conceptualization, Data curation, Formal analysis, Investigation, Methodology, Software, Validation, Visualization, Writing-original draft, Writing-review & editing. **Benjamin Marie:** Data curation, Formal analysis, Investigation, Software, Visualization, Writing-review & editing. **Soizic Morin**: Formal analysis, Investigation, Visualization, Writing-review & editing. **Pierre Foucault**: Software, Writing-review & editing. **Mathieu Milhe-Poutingon**: Formal analysis, Investigation. **Siann Chalvin**: Formal analysis, Investigation. **Clémentine Gelber**: Project administration, Resources, Writing-review & editing. **Patrick Balodni-Andrey**: Funding acquisition, Resources, Writing-review & editing. **Nicholas Gurieff**: Funding acquisition, Resources, Writing-review & editing. **Claude Fortin**: Supervision, Writing-review & editing. **Séverine Le Faucheur**: Conceptualization, Funding acquisition, Investigation, Project administration, Resources, Supervision, Validation, Writing-original draft, Writing-review & editing.

## Associated content

Detailed results on the physicochemical parameters of the exposure media; biofilm metal contents; Co effects on biofilm biomass, chlorophyll *a* content, diatom mortality and density; alpha- and beta-diversity indexes and compositions of prokaryotic and eukaryotic communities; untargeted and annotated meta-metabolomic fingerprints, T-*SNE* molecular network, BMDs and BMTs values; determination of CRIDeR/CRIDaR and TRIDeR/TRIDaR initiation thresholds, dissimilarity between biofilm meta-metabolomes, prokaryotic and eukaryotic communities are presented in supporting information.

## Acknowledgments

We would like to thank the members of the Environment & Sustainable Development Team, at Pôle d’Etudes et de Recherche de Lacq (PERL) for access to the TotalEnergies facilities for mesocosm exposures and assistance during experimentation. We are grateful to the UMR 7245 MCAM, Muséum National d’Histoire Naturelle, Paris, France, especially the “Cyanobactéries, Cyanotoxines et Environnement” (CCE) team and the Plateau technique de spectrométrie de masse bio-organique (PtSMB) for laboratories and technical facilities for meta-metabolomics analysis. Part of the experiments (amplicon sequencing) were performed at the PGTB (doi:10.15454/1.5572396583599417E12) with the assistance of Zachary Allouche and Zoé Delporte. We also thank Claire Gassie (Universite de Pau et des Pays de l’Adour, E2S-UPPA, CNRS, IPREM, Pau, France) for her help in characterization of prokaryotic and eukaryotic communities. Special thanks to Scott Hepditch (Institut National de la Recherche Scientifique – Eau Terre Environnement, Québec, Canada) for his language assistance.

## Abbreviations

AICc: second-order Akaike’s information criterion
BMD: benchmark-dose
BMT: benchmark-time
CA: cellulose acetate
CRIDaR: concentration range inducing damage responses
CRIDeR: concentration range inducing defense responses
DOC: dissolved organic carbon
ECDF: empirical cumulative density function
LC-HRMS: liquid chromatography-high resolution mass spectrometry
MOTA: multivariate omics trajectory analysis
PCoA: principal coordinate analysis
PP: polypropylene
TRIDaR: time range inducing damage responses
TRIDeR: time range inducing defense responses.

