## Supplementary Information Figures for "New sensitive tools to characterize meta-metabolome response to short- and long-term cobalt exposure in dynamic river biofilm communities"

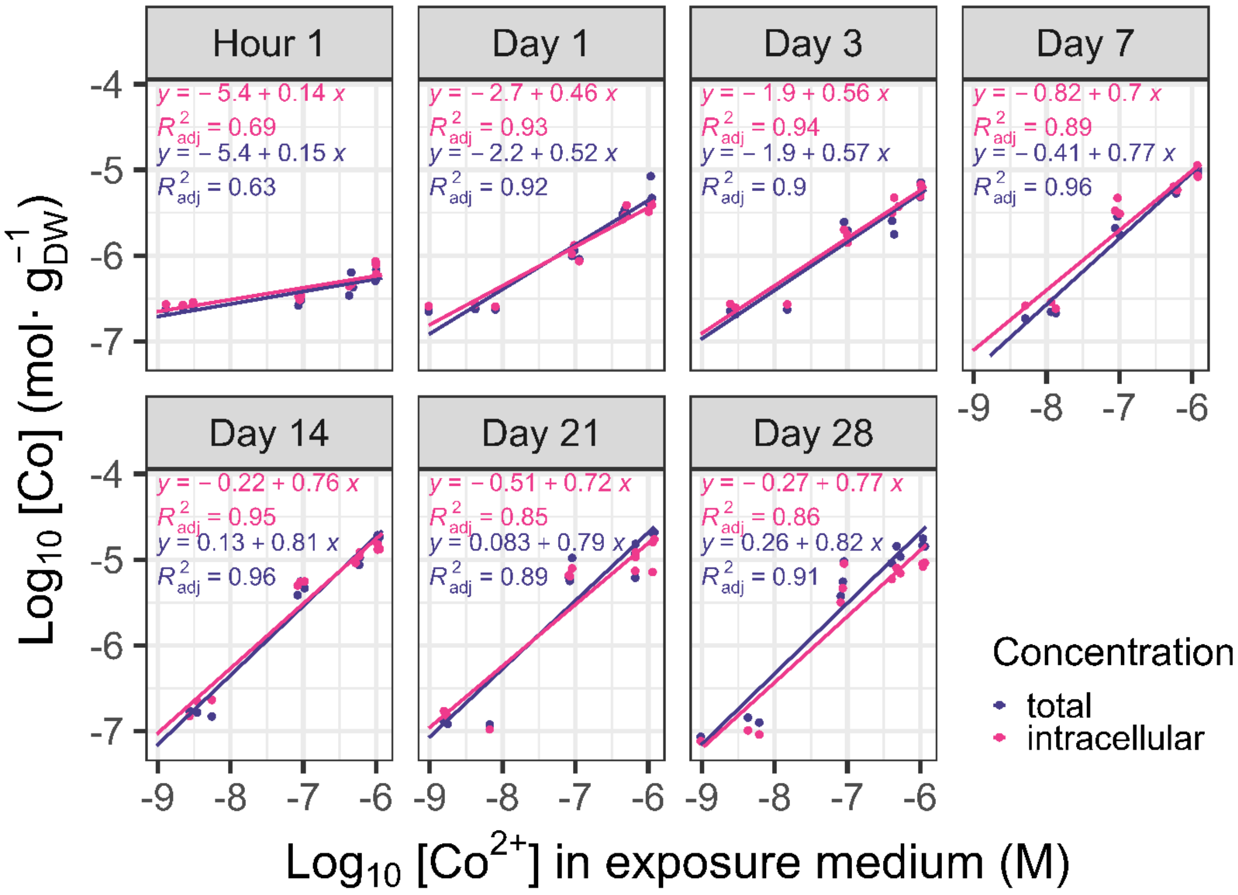


**Fig. A.1**: Cobalt bioaccumulation by biofilms. Blue and pink lines represent regressions of total and intracellular Co contents as a function of free Co concentration in exposure medium at different exposure times, respectively.

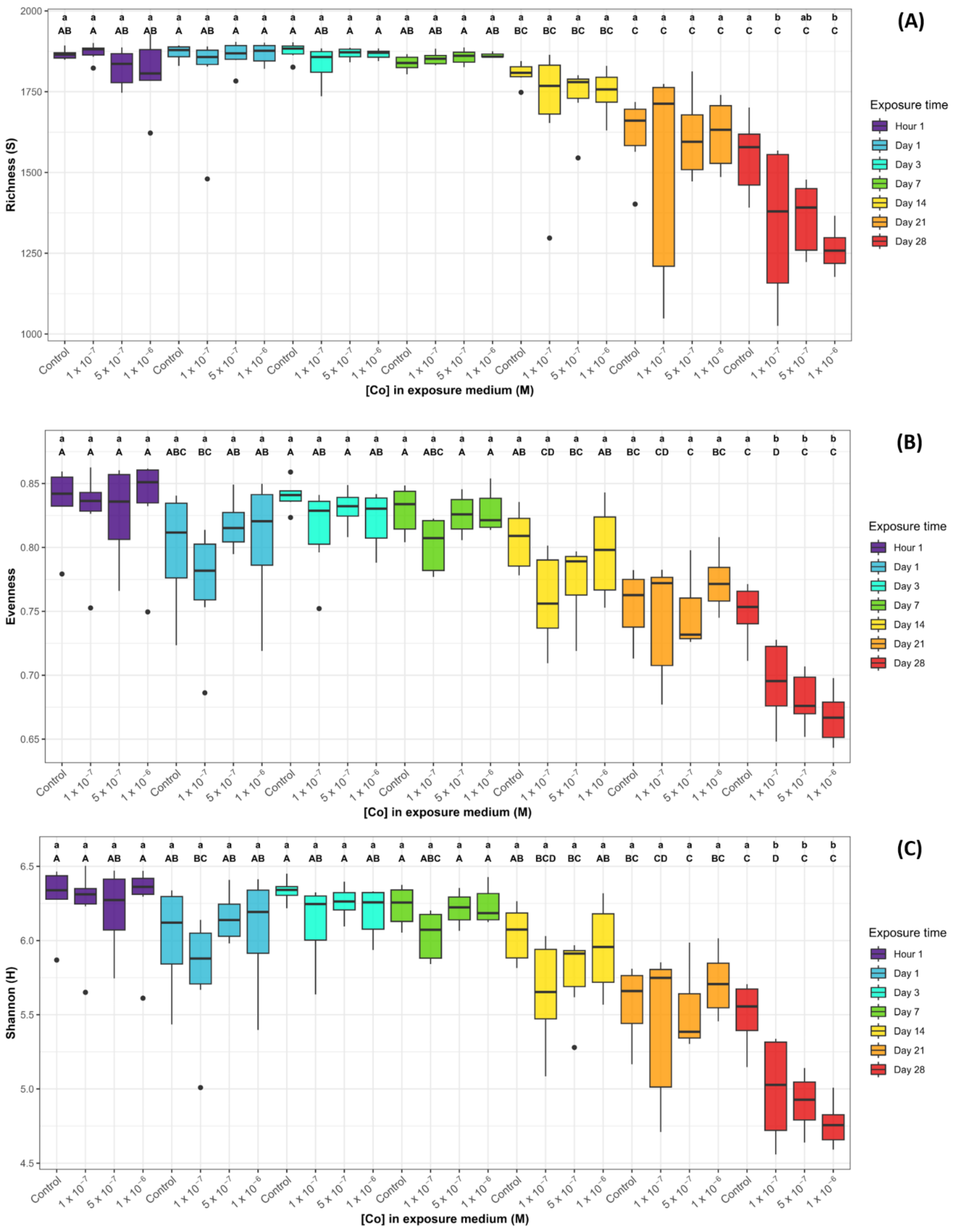


**Fig. A.2**: Prokaryotic (A) ASVs richness, (B) Pielou's evenness and (C) Shannon indices of biofilms exposed to different Co concentrations during 28 days. The lower-case letters correspond to the significative groups defined by a Dunn’s *post-hoc* test (*p-values* < 0.05) performed after Kruskal-Wallis non-parametric tests among Co exposure concentrations at each exposure time. Capital letters correspond to the significative groups defined by a Dunn’s *post-hoc* test (*p-values* < 0.05) performed after Kruskal-Wallis non-parametric tests among the exposure times for each exposure concentration.

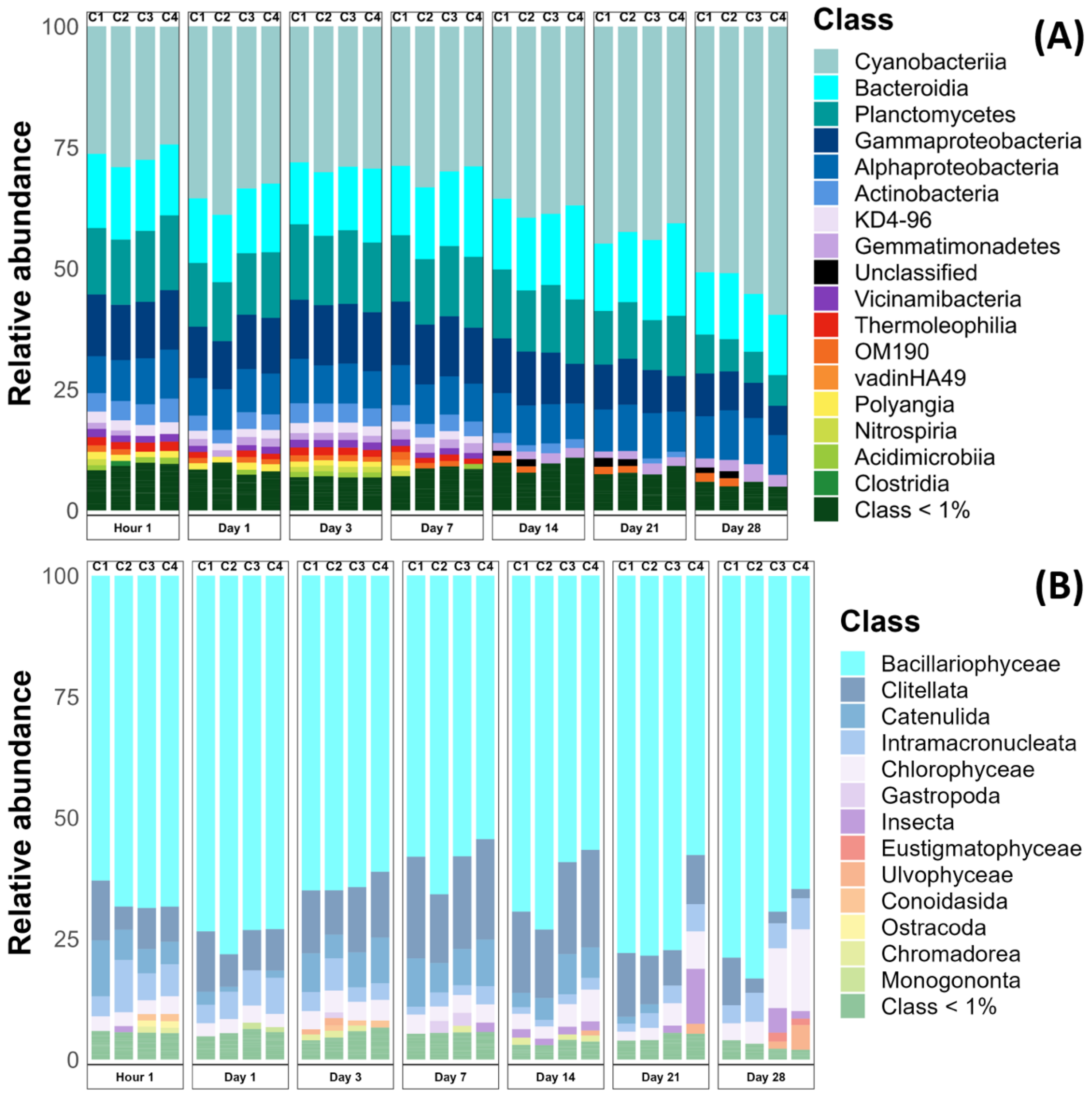


**Fig. A.3**: Co concentrations and exposure time effects on biofilm composition. (A) prokaryotic community composition at the class level (median values of samples belonging the same time step and exposure concentration group). (B) Eukaryotic community composition at the class level (median values of samples belonging the same time step and exposure concentration groupl; C1: control biofilms exposed at the background concentration, C2: biofilms exposed at $1 \times 10^{-7}$ M, C3: biofilms exposed at $5 \times 10^{-7}$ M and C4: biofilms exposed at $1 \times 10^{-6}$ M).

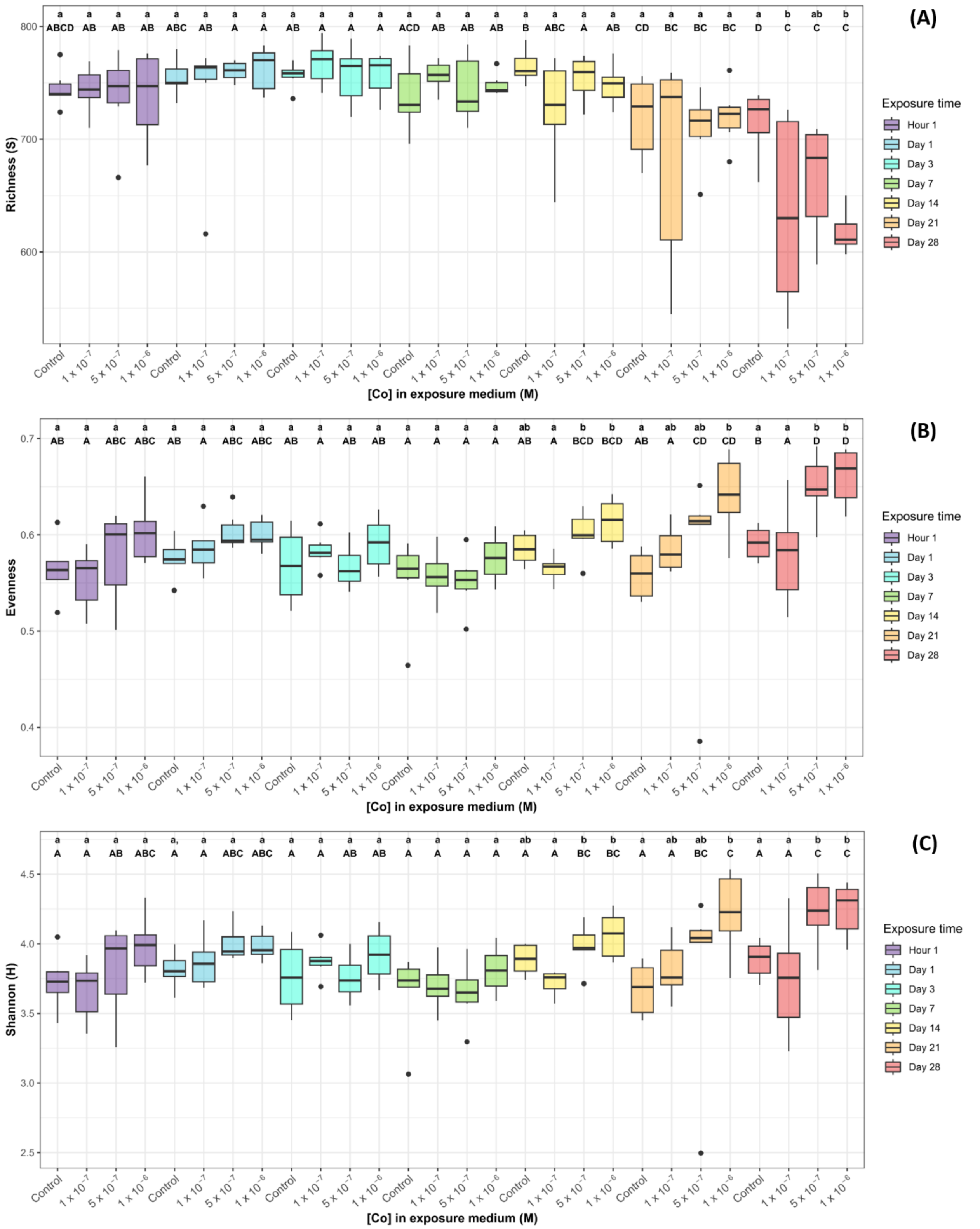


**Fig. A.4**: Eukaryote (A) ASVs richness, (B) Pielou's evenness and (C) Shannon indices of biofilms exposed to different Co concentrations during 28 days. The lower-case letters correspond to the significative groups defined by a Dunn's *post-hoc* test (*p-values* < 0.05) performed after Kruskal-Wallis non-parametric tests among Co exposure concentrations at each exposure time. Capital letters correspond to the significative groups defined by a Dunn's *post-hoc* test (*p-values* < 0.05) performed after Kruskal-Wallis non-parametric tests among the exposure times for each exposure concentration.

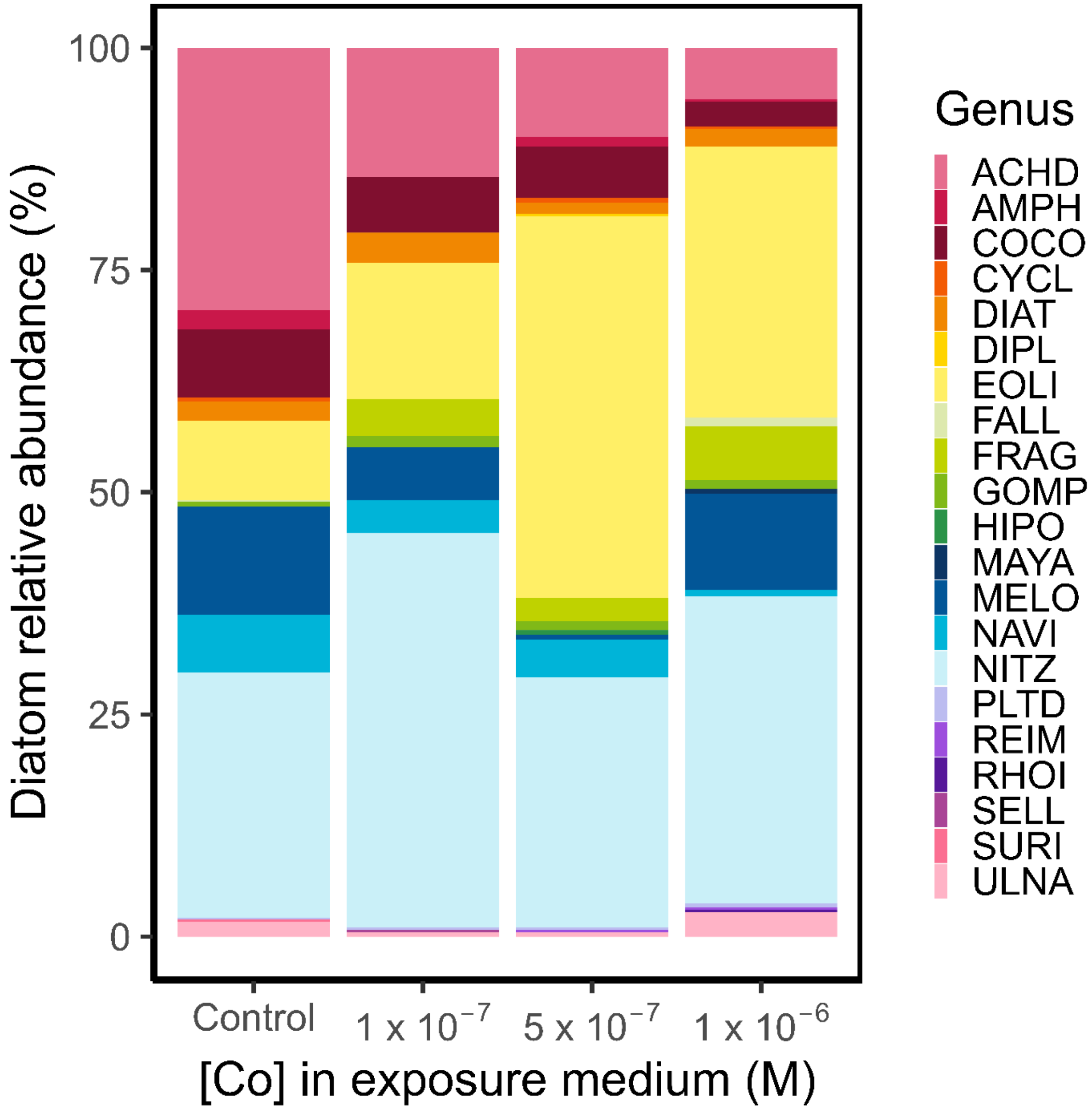


**Fig. A.5**: Diatom relative abundances (main genera) identified on day 28 in pooled replicate samples. Abbreviations for genera are: ACHD: *Achnanthidium*, AMPH: *Amphora*, COCO: *Cocconeis*, CYCL: *Cyclotella*, DIAT: *Diatoma*, DIPL: *Diploneis*, EOLI: *Eolimna*, FALL: *Fallacia*, FRAG: *Fragilaria*, GOMP: *Gomphonema*, HIPO: *Hippodonta*, MAYA: *Mayamaea*, MELO: *Melosira*, NAVI: *Navicula*, NITZ: *Nitzschia*, PLTD: *Planothidium*, REIM: *Reimeria*, RHOI: *Rhoicosphenia*, SELL: *Sellaphora*, SURI: *Surirella*, ULNA: *Ulnaria*.

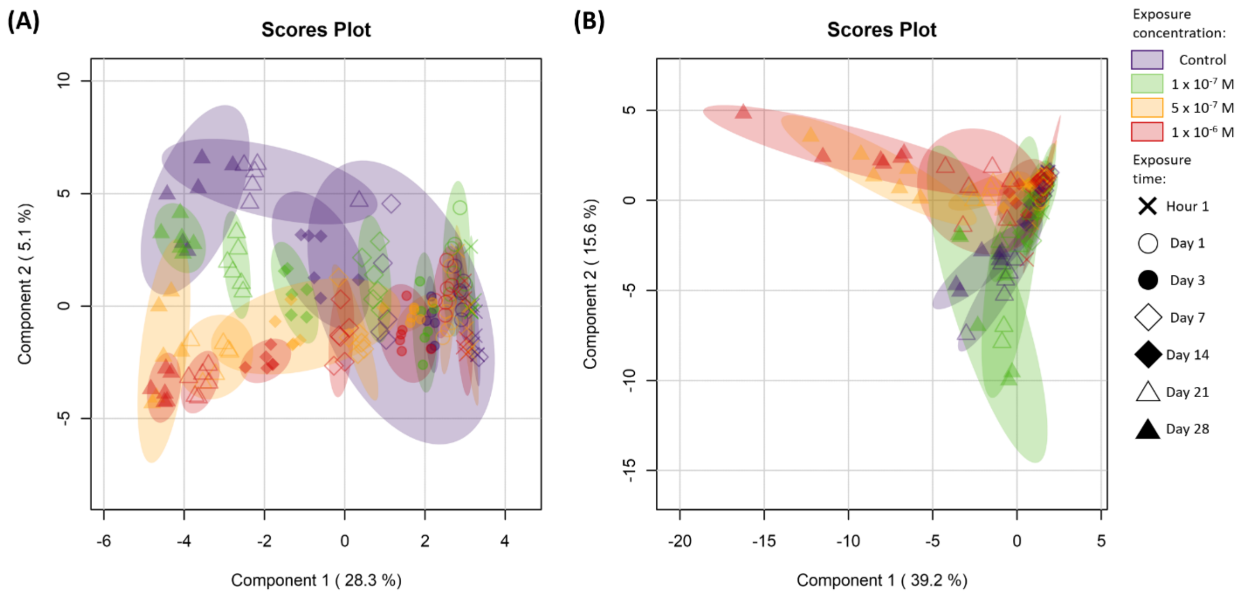


**Fig. A.6**: Individual score plot generated from sPLS-DA analysis performed with 10 variables to components 1-2, (A) for untargeted meta-metabolomic features and (B) for annotated metabolites.

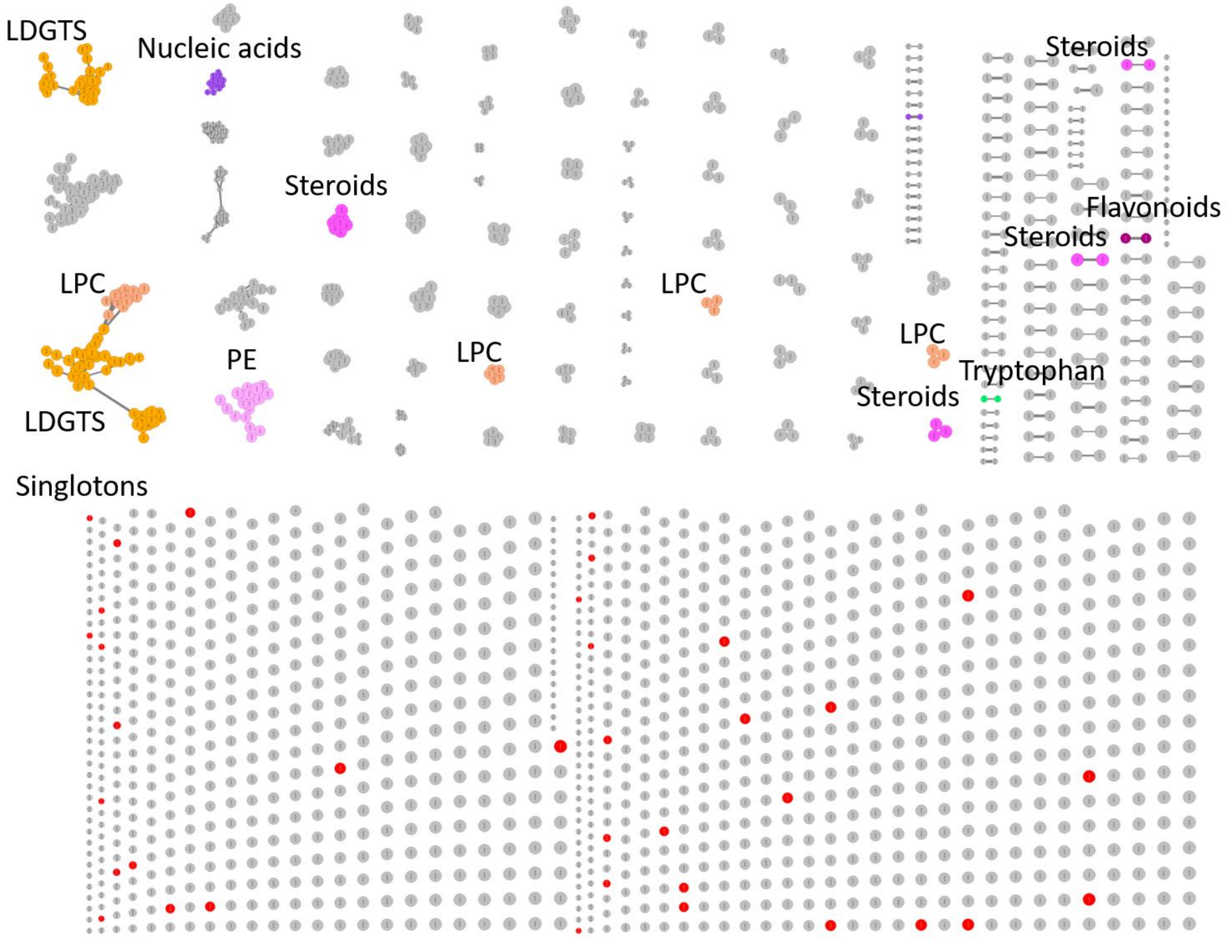


**Fig. A.7**: T-*SNE* molecular network generated with Metgem 1.3.6 based on LC-MSMS dataset obtained for all samples. The various colors show the different molecular families that constitute the different annotated clusters according to molecular fragment similarities with standards available in public database (GNPS, HMDB, NIH and EMBL).

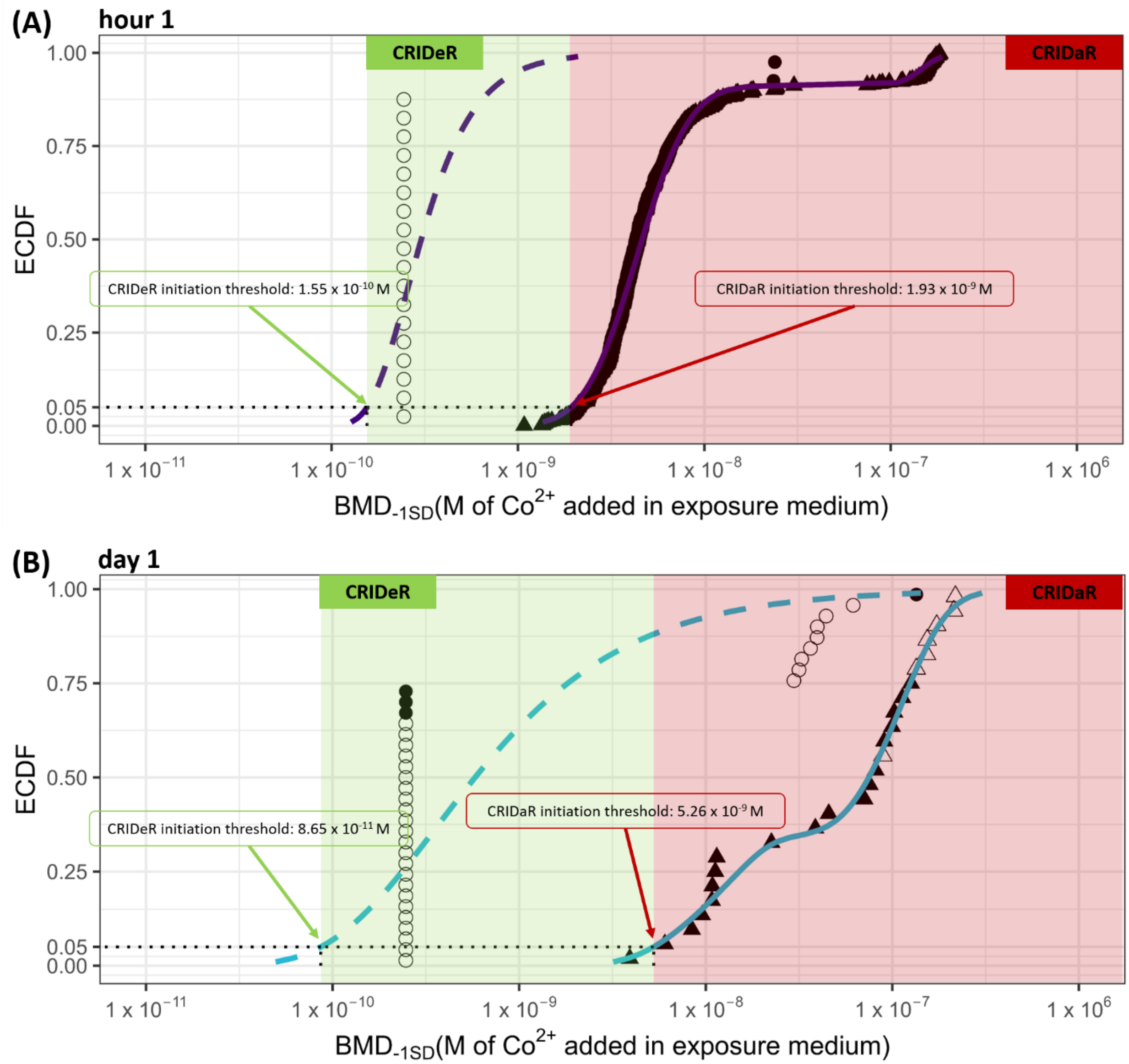

(A)
hour 1
CRIDeR
CRIDaR
ECDF
1.00
0.75
0.50
0.25
0.05
0.00
CRIDeR initiation threshold: 1.55 x 10-10 M
CRIDaR initiation threshold: 1.93 x 10-9 M
1 x 10−11
1 x 10−10
1 x 10−9
1 x 10−8
1 x 10−7
1 x 106
BMD-1SD(M of Co2+ added in exposure medium)
(B)
day 1
CRIDeR
CRIDaR
ECDF
1.00
0.75
0.50
0.25
0.05
0.00
CRIDaR initiation threshold: 5.26 x 10-9 M
CRIDeR initiation threshold: 8.65 x 10-11 M
1 x 10−11
1 x 10−10
1 x 10−9
1 x 10−8
1 x 10−7
1 x 106
BMD-1SD(M of Co2+ added in exposure medium)

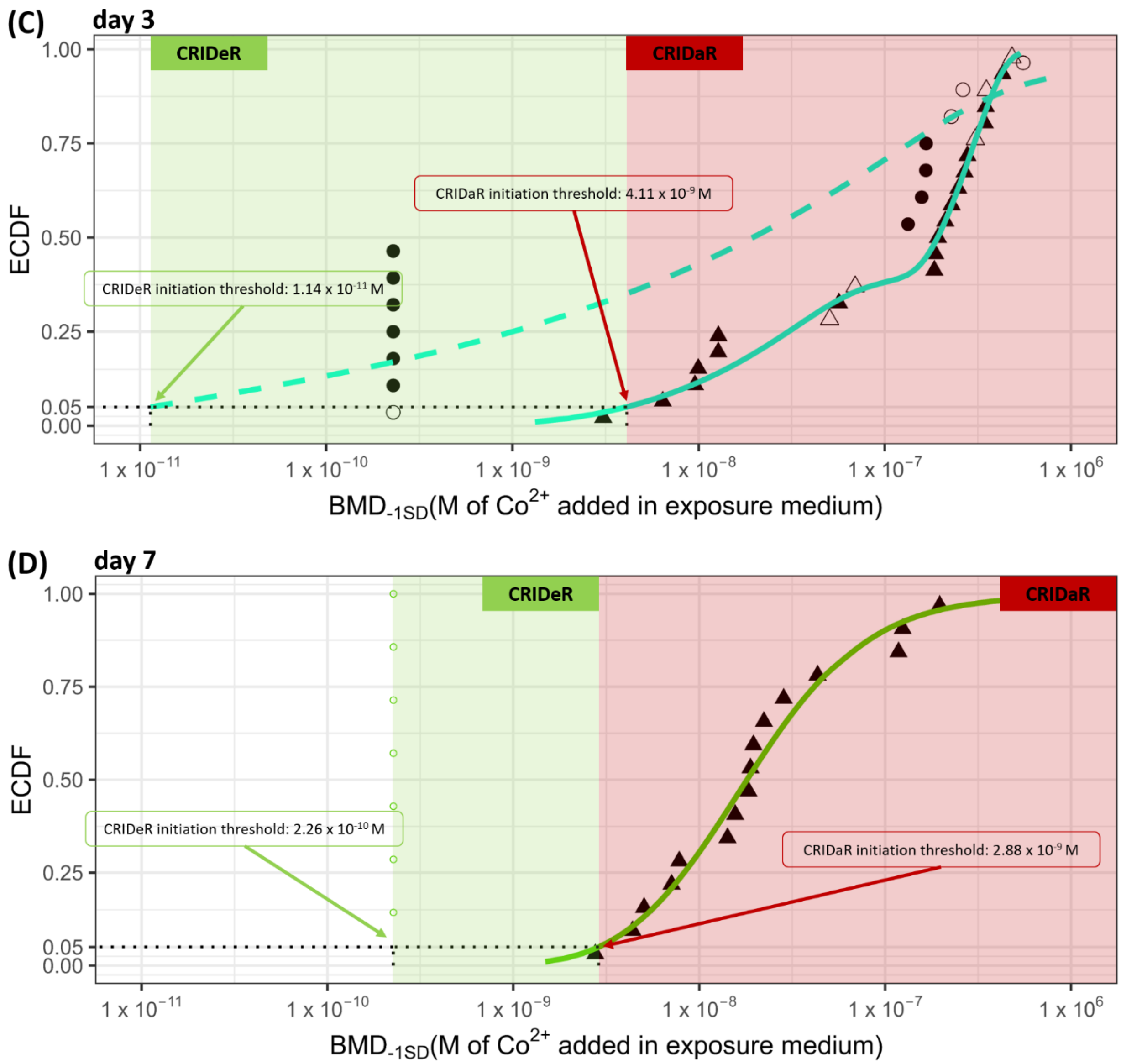
(C)
day 3
CRIDeR
CRIDaR
CRIDaR initiation threshold: 4.11 x 10^-9 M
CRIDeR initiation threshold: 1.14 x 10^-11 M
ECDF
1.00
0.75
0.50
0.25
0.05
0.00
1 x 10^−11
1 x 10^−10
1 x 10^−9
1 x 10^−8
1 x 10^−7
1 x 10^6
BMD-1SD(M of Co2+ added in exposure medium)
(D)
day 7
CRIDeR
CRIDaR
CRIDeR initiation threshold: 2.26 x 10^-10 M
CRIDaR initiation threshold: 2.88 x 10^-9 M
ECDF
1.00
0.75
0.50
0.25
0.05
0.00
1 x 10^−11
1 x 10^−10
1 x 10^−9
1 x 10^−8
1 x 10^−7
1 x 10^6
BMD-1SD(M of Co2+ added in exposure medium)

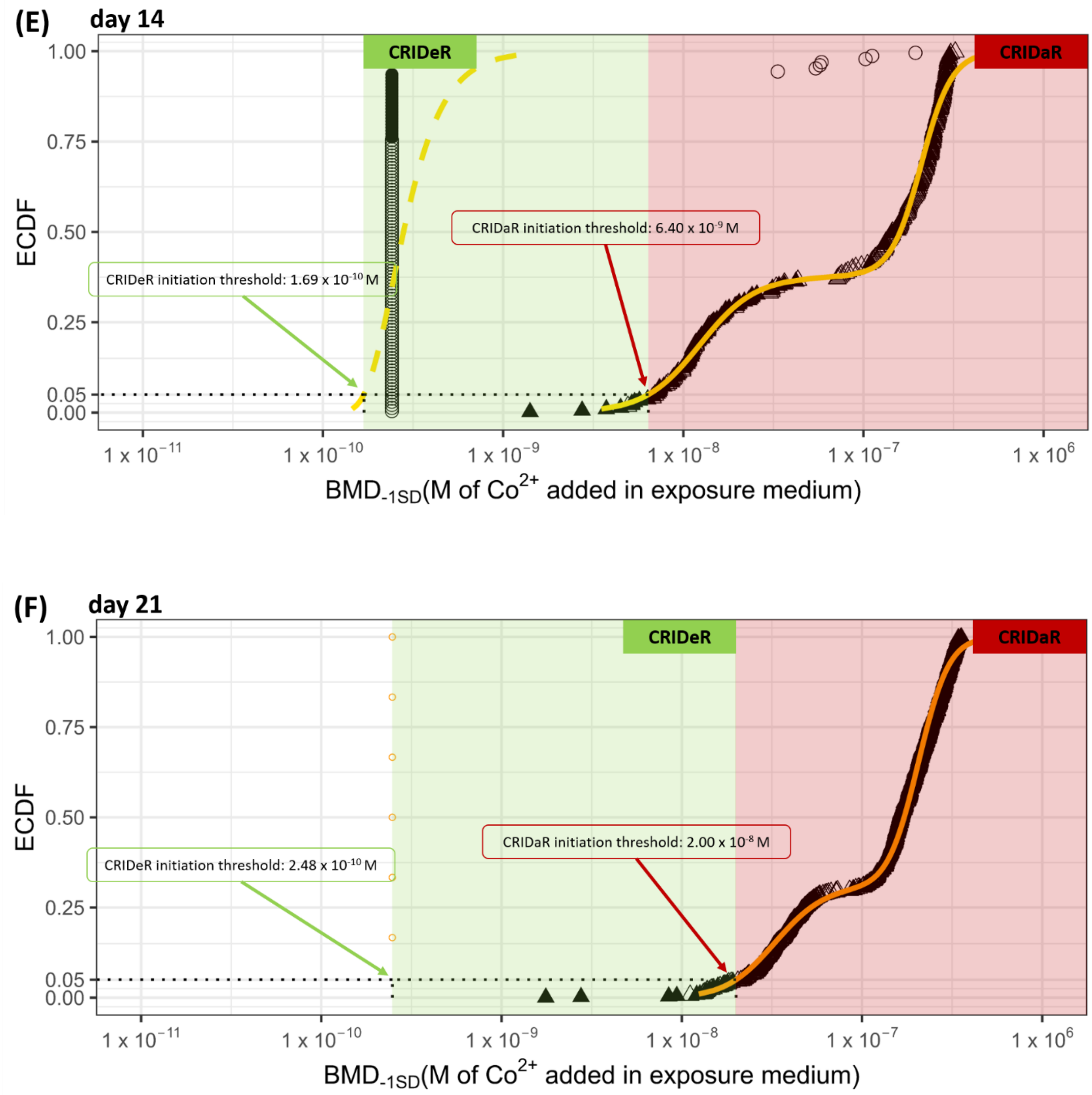
(E)
day 14
CRIDeR
CRIDaR
CRIDaR initiation threshold: 6.40 x 10-9 M
CRIDeR initiation threshold: 1.69 x 10-10 M
ECDF
1.00
0.75
0.50
0.25
0.05
0.00
1 x 10−11
1 x 10−10
1 x 10−9
1 x 10−8
1 x 10−7
1 x 106
BMD-1SD(M of Co2+ added in exposure medium)
(F)
day 21
CRIDeR
CRIDaR
CRIDaR initiation threshold: 2.00 x 10-8 M
CRIDeR initiation threshold: 2.48 x 10-10 M
ECDF
1.00
0.75
0.50
0.25
0.05
0.00
1 x 10−11
1 x 10−10
1 x 10−9
1 x 10−8
1 x 10−7
1 x 106
BMD-1SD(M of Co2+ added in exposure medium)

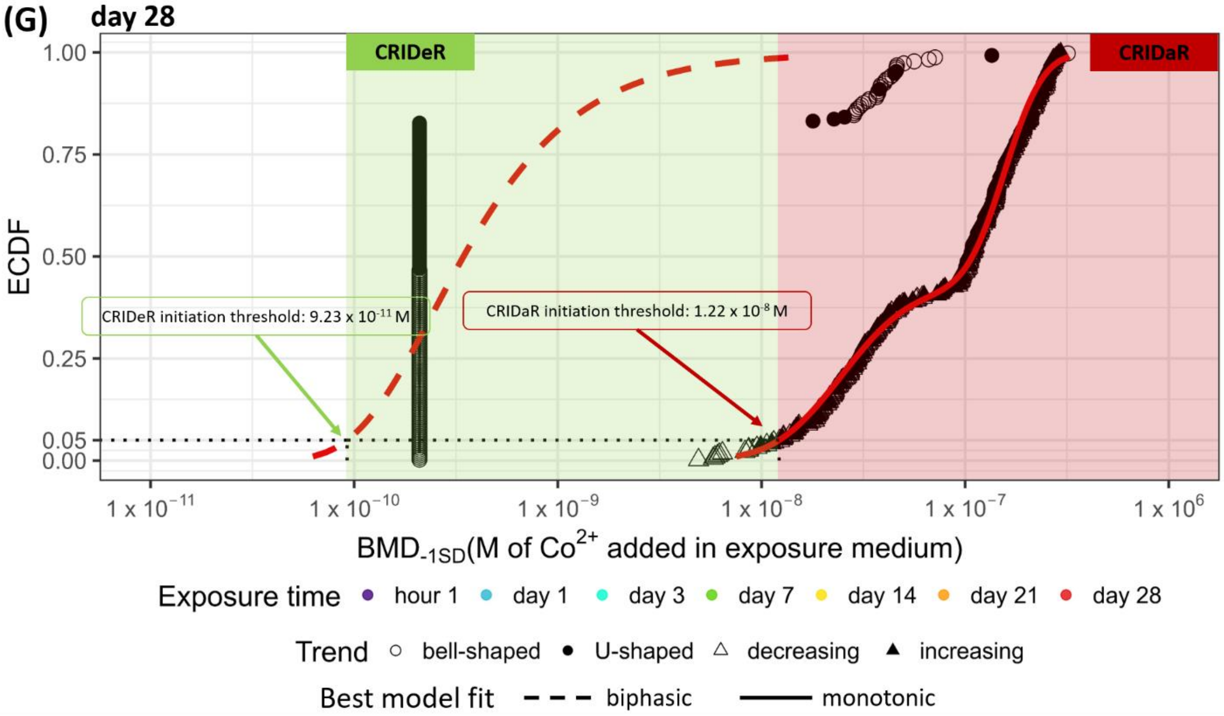


**Fig. A.8**: Determination of CRIDeR and CRIDaR initiation thresholds for each time step from the best fit model for ECDFs of untargeted meta-metabolomic features with biphasic (bell- or U-shaped) and monotonic (decreasing or increasing) trends, respectively. The CRIDeR and CRIDaR initiation thresholds are the hazardous concentration at which 5% of untargeted meta-metabolomic features will potentially be affected on the basis of their BMDs. (A) CRIDeR and CRIDar at hour 1; (B) CRIDeR and CRIDaR at day 1; (C) CRIDeR and CRIDaR at day 3; (D) CRIDeR and CRIDaR at day 7; (E) CRIDeR and CRIDaR at day 14; (F) CRIDeR and CRIDaR at day 21; (G) CRIDeR and CRIDaR at day 28. * CRIDeR threshold determined directly with the BMD of the first untargeted meta-metabolomic feature impacted because all distributions failed to fit.

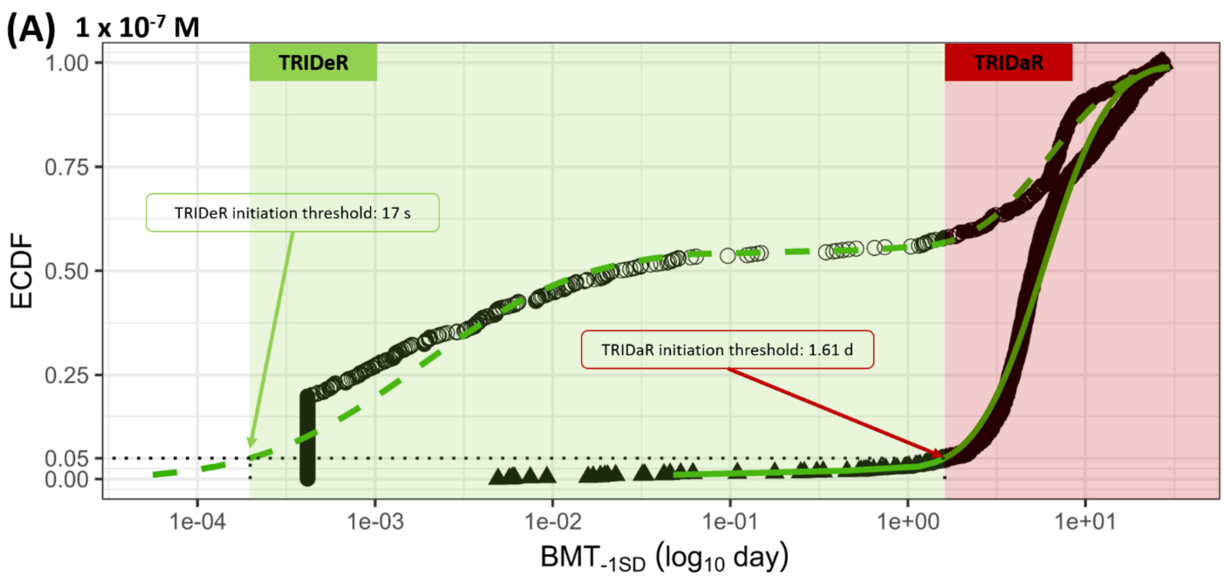

(A) 1 x 10-7 M
TRIDeR
TRIDaR
TRIDeR initiation threshold: 17 s
TRIDaR initiation threshold: 1.61 d
ECDF
1.00
0.75
0.50
0.25
0.05
0.00
1e-04
1e-03
1e-02
1e-01
1e+00
1e+01
BMT-1SD (log10 day)


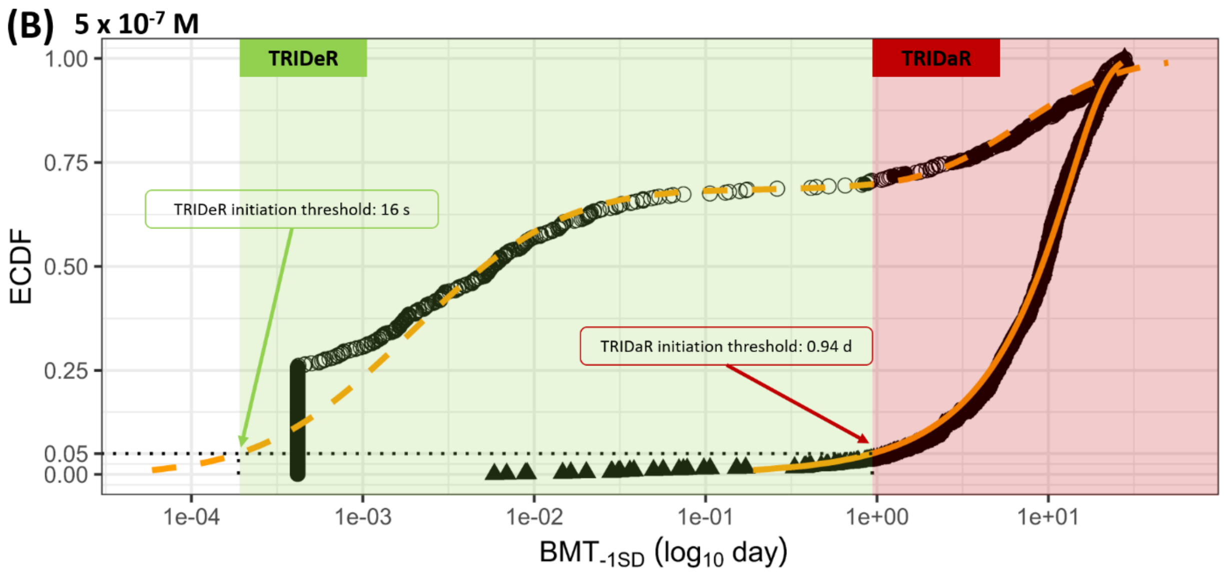

(B) 5 x 10-7 M
TRIDeR
TRIDaR
TRIDeR initiation threshold: 16 s
TRIDaR initiation threshold: 0.94 d
ECDF
1.00
0.75
0.50
0.25
0.05
0.00
1e-04
1e-03
1e-02
1e-01
1e+00
1e+01
BMT-1SD (log10 day)

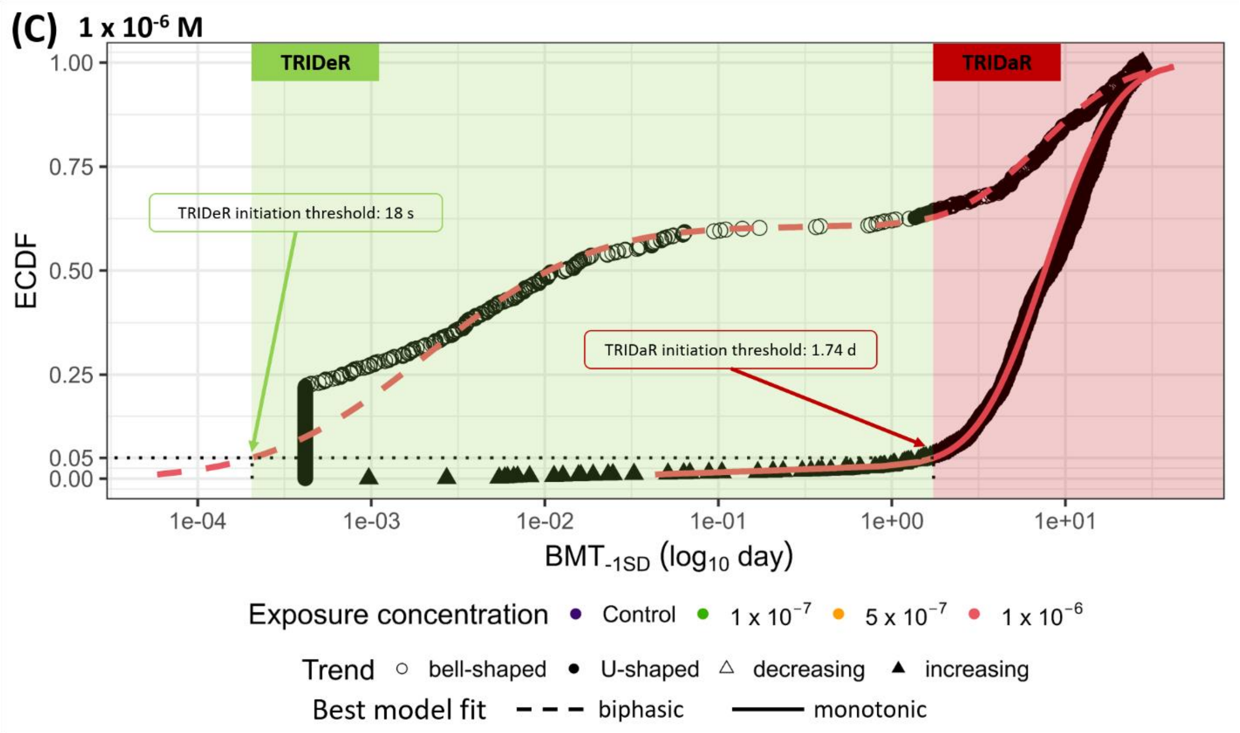


**Fig. A.9**: Determination of TRIDeR and TRIDaR initiation thresholds for each exposure concentration from the best fit model for ECDFs of untargeted meta-metabolomic features with biphasic (bell- or U-shaped) and monotonic (decreasing or increasing) trends, respectively. The TRIDeR and TRIDaR initiation thresholds are the hazardous exposure time at which 5% of untargeted meta-metabolomic features will potentially be affected on the basis of their BMTs. (A) TRIDeR and TRIDaR at 1 x $10^{-7}$ M; (B) TRIDeR and TRIDaR at 5 x $10^{-7}$ M; (C) TRIDeR and TRIDaR at at 1 x $10^{-6}$ M.

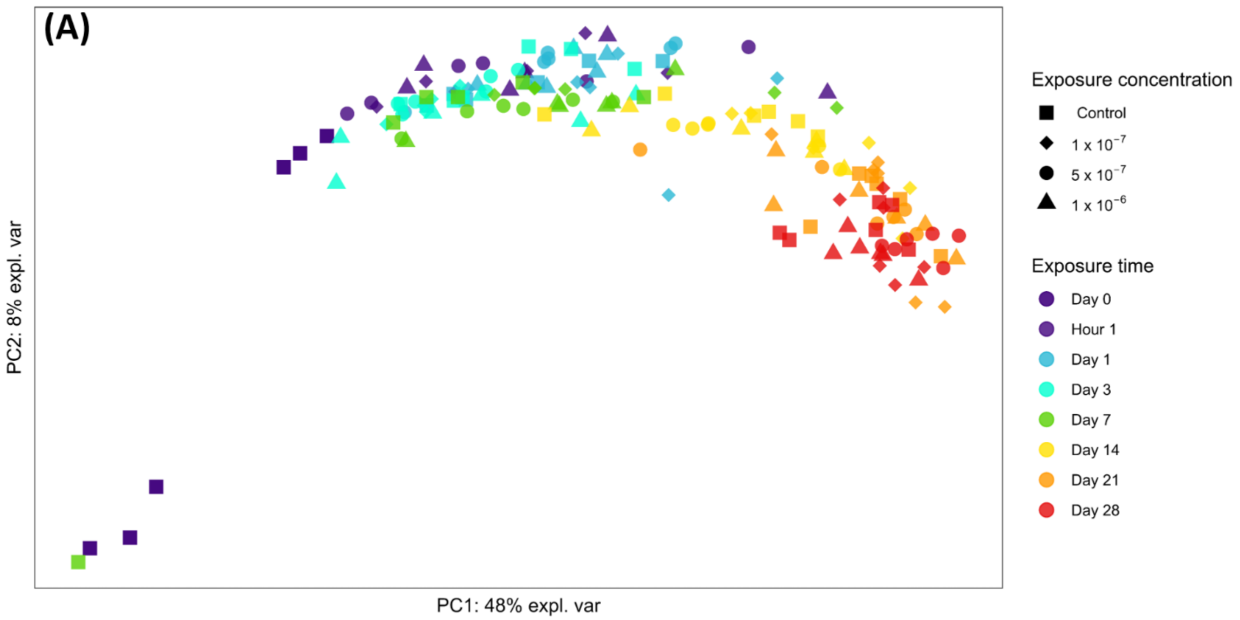
(A)
PC2: 8% expl. var
PC1: 48% expl. var
Exposure concentration
Control
1 x 10$^{-7}$
5 x 10$^{-7}$
1 x 10$^{-6}$
Exposure time
Day 0
Hour 1
Day 1
Day 3
Day 7
Day 14
Day 21
Day 28

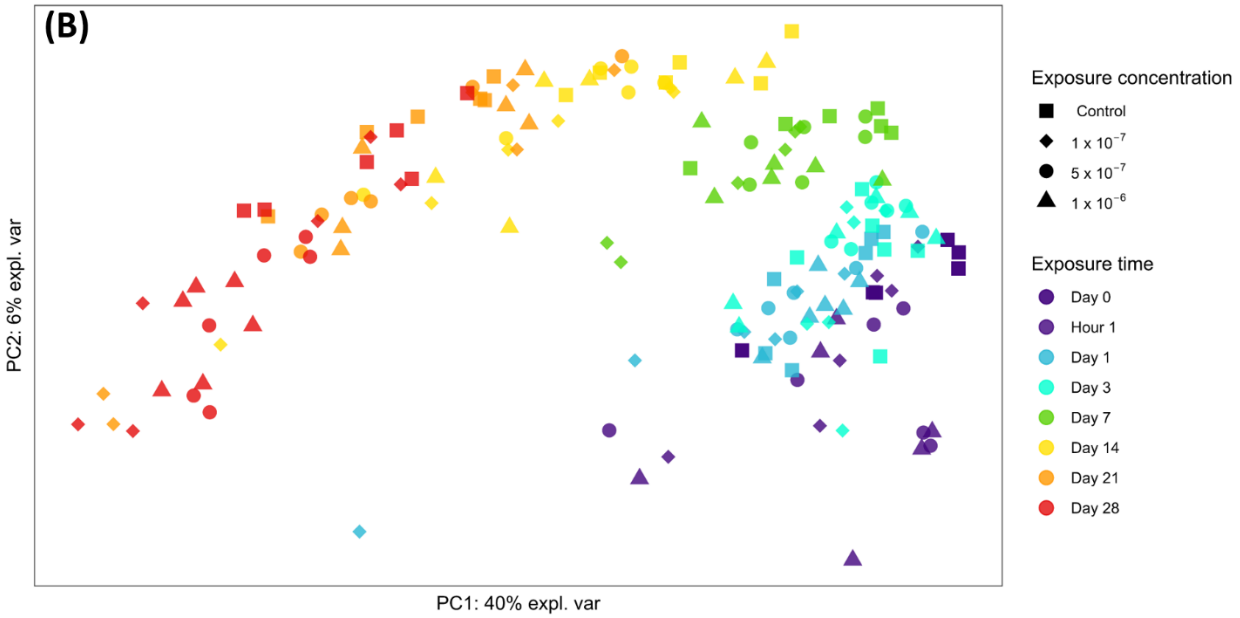
(B)
PC2: 6% expl. var
PC1: 40% expl. var
Exposure concentration
Control
1 x 10$^{-7}$
5 x 10$^{-7}$
1 x 10$^{-6}$
Exposure time
Day 0
Hour 1
Day 1
Day 3
Day 7
Day 14
Day 21
Day 28

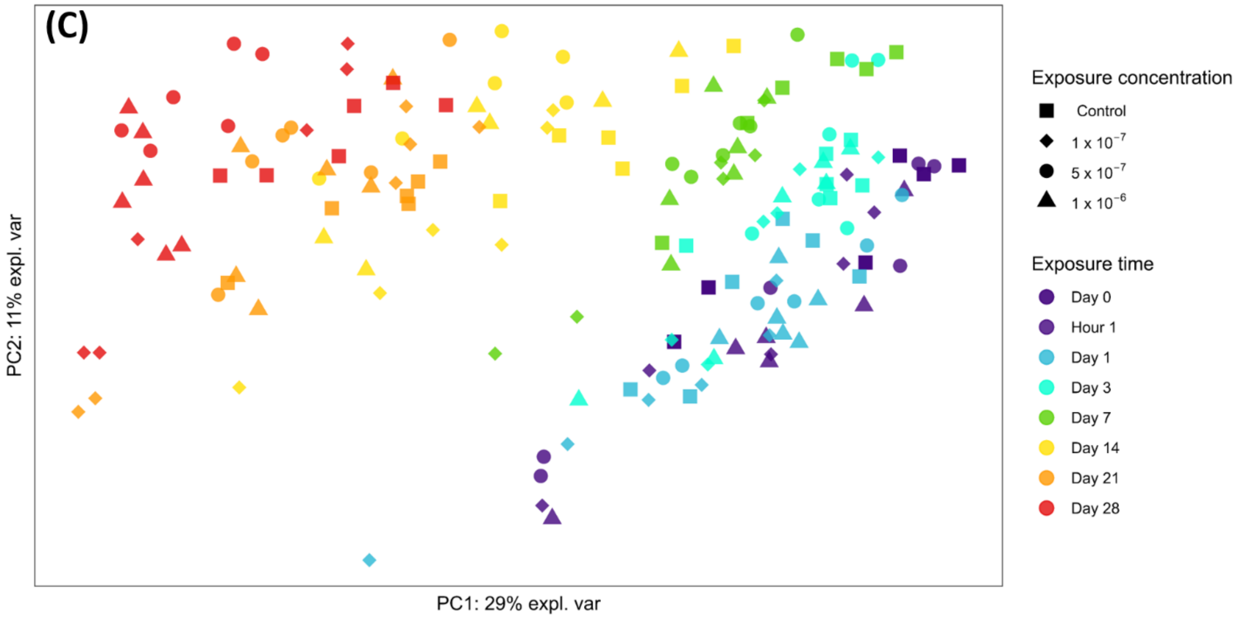


**Fig. A.10**: Dissimilarity between biofilm meta-metabolomes, prokaryotic and eukaryotic communities as a function of Co concentration and exposure time. (A) Principal component analysis of the meta-metabolome composition of exposed and control biofilms over time used for MOTA. (B) Principal component analysis of the prokaryotic community of exposed and unexposed biofilms over time used for MOTA. (C) Principal component analysis of the eukaryotic community of exposed and unexposed biofilms over time used for MOTA.

**Tables**

See .xlsx file

**Table A.1**: Physicochemical parameters of the exposure media during 28 days of exposure. The lower-case letters correspond to the significative groups defined by a Dunn's *post-hoc* test (*p-values* < 0.05) performed after Kruskal-Wallis non-parametric tests among Co exposure concentrations at each exposure time. Capital letters correspond to the significative groups defined by a Dunn's *post-hoc* test (*p-values* < 0.05) performed after Kruskal-Wallis non-parametric tests among the exposure times for each exposure concentration.

**Table A.2**: Total and intracellular Co contents bioaccumulated by biofilms during 28 days of exposure. The lower-case letters correspond to the significative groups defined by a Dunn's *post-hoc* test (*p-values* < 0.05) performed after Kruskal-Wallis non-parametric tests among Co exposure concentrations at each exposure time. Capital letters correspond to the significative groups defined by a Dunn's *post-hoc* test (*p-values* < 0.05) performed after Kruskal-Wallis non-parametric tests among the exposure times for each exposure concentration.

**Table A.3**: Intracellular metal contents bioaccumulated by biofilms during 28 days of exposure. The lower-case letters correspond to the significative groups defined by a Dunn's *post-hoc* test (*p-values* < 0.05) performed after Kruskal-Wallis non-parametric tests among Co exposure concentrations at each exposure time. Capital letters correspond to the significative groups defined by a Dunn's *post-hoc* test (*p-values* < 0.05) performed after Kruskal-Wallis non-parametric tests among the exposure times for each exposure concentration.

**Table A.4**: Cobalt effects on biofilm biomass, chlorophyll *a* content, diatom mortality and diatom density during 28 days of exposure. The lower-case letters correspond to the significative groups defined by a Dunn's *post-hoc* test (*p-values* < 0.05) performed after Kruskal-Wallis non-parametric tests among Co exposure concentrations at each exposure time. Capital letters correspond to the significative groups defined by a Dunn's *post-hoc* test (*p-values* < 0.05) performed after Kruskal-Wallis non-parametric tests among the exposure times for each exposure concentration.

**Table A.5**: Cobalt effect on beta diversity metrics of prokaryotic communities during 28 days of exposure, group comparison scores and significance levels (exposure concentration groups: C1: control biofilms exposed at the background concentration, C2: biofilms exposed at 1 x $10^{-7}$ M, C3: biofilms exposed at 5 x $10^{-7}$ M and C4: biofilms exposed at 1 x $10^{-6}$ M).

**Table A.6**: Cobalt effect on beta diversity metrics of eukaryotic communities during 28 days of exposure, group comparison scores and significance levels (exposure concentration groups: C1: control biofilms exposed at the background concentration, C2: biofilms exposed at 1 x $10^{-7}$ M, C3: biofilms exposed at 5 x $10^{-7}$ M and C4: biofilms exposed at 1 x $10^{-6}$ M.

**Table A.7**: Cobalt effect on diatom species relative abundances identified on day 28 in pooled replicate samples.

**Table A.8**: Cobalt effect on biofilm meta-metabolome during 28 days of exposure, group comparison scores and significance levels (exposure concentration groups: C1: control biofilms exposed at the background concentration, C2: biofilms exposed at $1 \times 10^{-7}$ M, C3: biofilms exposed at $5 \times 10^{-7}$ M and C4: biofilms exposed at $1 \times 10^{-6}$ M).

**Table A.9**: Cobalt effect on biofilm annotated metabolites during 28 days of exposure, group comparison scores and significance levels (exposure concentration groups: C1: control biofilms exposed at the background concentration, C2: biofilms exposed at $1 \times 10^{-7}$ M, C3: biofilms exposed at $5 \times 10^{-7}$ M and C4: biofilms exposed at $1 \times 10^{-6}$ M).

**Table A.10**: Proportions of untargeted meta-metabolomic features whose response is significantly impacted by Co concentrations according to their dose-response model curve trend and BMD value.

**Table A.11**: Goodness of model fit for the CRIDeR and CRIDaR initiation threshold determinations, in bold the best predictive model (with the lowest AICc).

**Table A.12**: Proportions of untargeted meta-metabolomic features whose response is significantly impacted by exposure time according to their dose-response model curve trend and BMT value.

**Table A.13**: Goodness of model fit for the TRIDeR and TRIDaR initiation threshold determinations, in bold the best predictive model (with the lowest AICc).
